# The nucleus follows an internal cellular scale during polarized root hair cell development

**DOI:** 10.1101/2025.08.21.671641

**Authors:** Jessica M. Orr, M. Arif Ashraf

## Abstract

The root hair cell is a product of asymmetric cell division, which grows in a polar manner, and thus is an attractive model cell type from a cellular biology aspect. Beyond the fundamental cell biology context, the root hair is involved in water and nutrient acquisition, making it important for agronomic applications. Nuclear positioning in the cell is crucial during root hair development. Often, textbooks demonstrate illustrations of the nucleus located at a fixed position from the tip of a root hair or at the very end of a root hair. The fundamental question is whether the nucleus follows a cellular scale during polarized growth. Maintaining the scale is a rudimentary biological process during development at the organismal and cellular levels. In this study, we altered root hairs through hormonal, nutrient, and environmental factors to decipher the cellular scale (no scale, universal scale, or internal scale) maintained by the nucleus. We utilized the live cell imaging combined with a quantitative cell biology approach and, surprisingly discovered that the nucleus always follows an internal scale in the root hair cell. This finding dramatically shifted our view about the nuclear position in a polarized cell and will have a potential to test across the tree of life. Altogether, understanding the cellular scale involved with nucleus positioning will have broad implications. It encourages a reexamination of textbooks and reinforces the agricultural importance in a changing climate.

## Introduction

The nucleus is a key cellular organelle for eukaryotic organisms. The very first observation of the nucleus at the cellular level can be traced to Robert Brown, who examined ∼4000 plant species and observed the epidermal cells under the simple light microscope (Ford, 1982). Soon, it was realized by Schwann, inspired by his lab mate and botanist Schleiden, that the nucleus is present beyond plant cells, and he observed a similar structure in cartilage cells (Wolpert, 1996). The nucleus is often illustrated as an organelle sitting at the center of the cell; however, it is a highly dynamic organelle, and its position varies depending on cell types and function within the cell (Gundersen and Worman, 2013). Dynamic nuclear movement is observed prior to asymmetric cell division and polarized cell growth (Ashraf and Facette, 2020; Ashraf et al., 2023; Hazelwood et al., 2025b; Hazelwood et al., 2025c). Plants serve as an excellent model system to study nuclear movement at the cellular, molecular, and developmental levels (Hazelwood et al., 2025b; Hazelwood et al., 2025c).

The lack of motility in plants is compensated for by polarized cell growth, which occurs in the root hair. Polarized growth of the root hair helps the below-ground root to reach water, minerals, and interact with soil and microbes (Zarebanadkouki et al., 2019). Root hair cells are a product of asymmetric cell division of epidermal cells of the model plant *Arabidopsis thaliana (A. thaliana)* (Kim and Dolan, 2011). In *A. thaliana* roots, root hair cells are arranged next to each other along the longitudinal axis, and separated by non-root hair cell files along the radial axis (Salazar-Henao et al., 2016). This unique patterning occurs due to asymmetric cell division of the epidermal cells, which results in trichoblasts (root hair cell) and atrichoblasts (non-root hair cell) (Fig. 1A). While non-root hair cells elongate longitudinally, root hair cells are shorter in length and develop a characteristic bulge (Fig. 1A). Following the bulge formation, the root hair cell grows polarly, and the nucleus moves directionally as well (Fig. 1A). Understanding nuclear positioning during polarized growth is fundamentally important for cellular organization and developmental patterning aspects.

**Figure 1:**
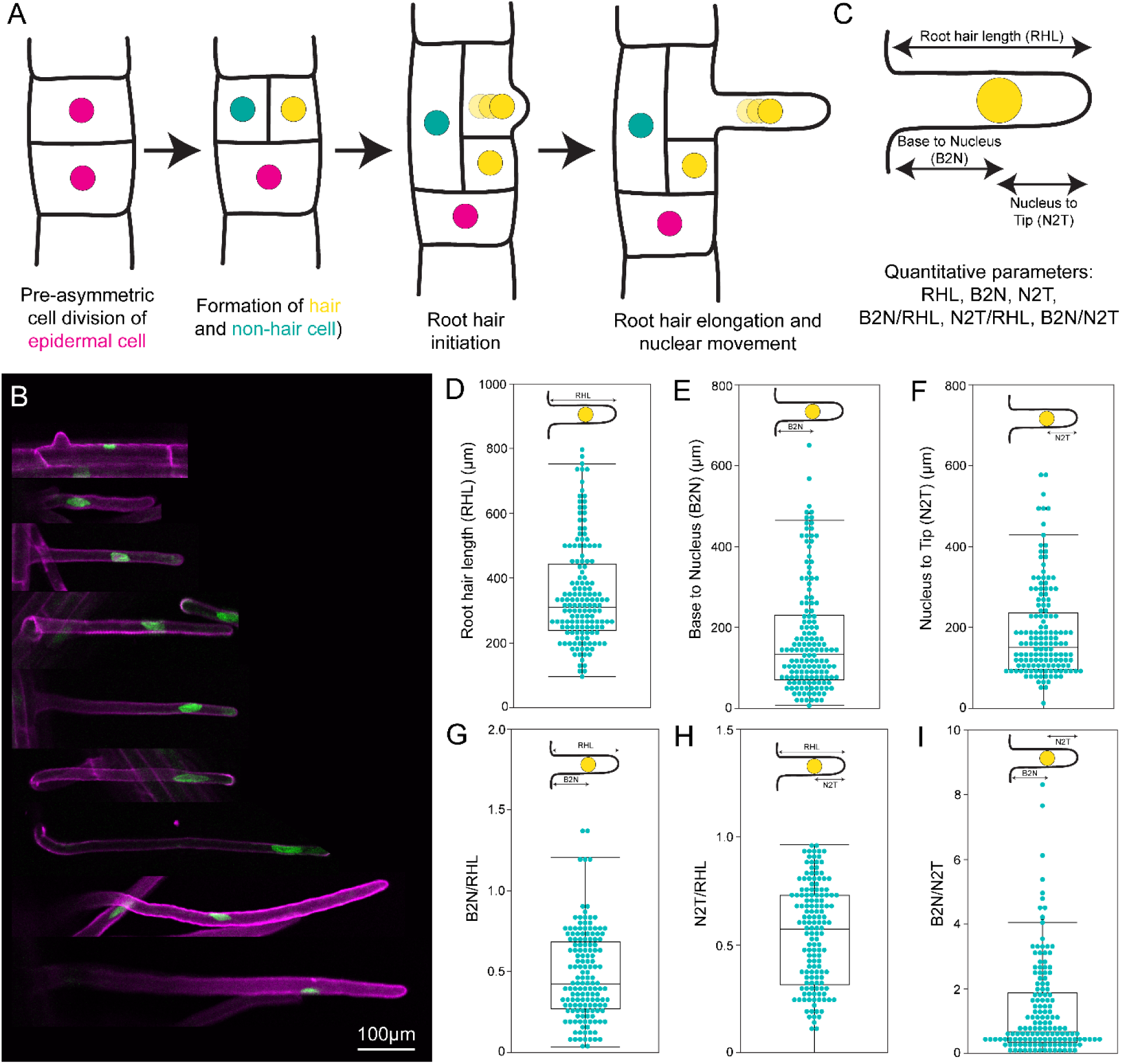
Nuclear positioning in root hair cell. (A) Model of root hair development in *A. thaliana*. Asymmetric cell division occurs, forming a hair cell and a non-hair cell. The non-hair cell elongates longitudinally, and the hair cell elongates polarly. The nucleus progresses along the length of the root hair cell. (B) Live cell imaging of root hair and nuclear position for wild-type. Scale = 100 µm. (C) Model showing the quantitative parameters measured from live cell imaging highlighted in B. (D-I) Quantification of root hair length (RHL) (D), quantification of the distance from the base to the nucleus (B2N) of the root hair (E), quantification of the distance from the nucleus to the tip (N2T) of the root hair (F), quantification of the ratio of B2N/RHL (G), quantification of the ratio of N2T/RHL (H) and quantification of the ratio of B2N/N2T (I). (D-I) The quantification area is shown at the top of each box plot. Each cyan dot represents a separate root hair that was measured.

Quantitative cell and developmental biology studies made major progress in understanding the root hair tip growth dynamics, nuclear speed, nuclear shape during polarized movement, and involvement of the cytoskeleton during this process using chemical and genetic approaches. For instance, early and late stages of root hair growth rates are 0.3 to 0.5 µm/min and 1.1 to 2.5 µm/min, respectively (Dolan et al., 1994). Within the root hair, nucleus movement varies between 0.3 to 2.0 µm/min (Chytilova et al., 2000). Moreover, the nuclear movement within the polarized root hair cell is identified as an actin-dependent process, not including microtubules (Chytilova et al., 2000; Ketelaar et al., 2002). A persistent issue with studying the root hair was the long-term imaging, which was resolved by the introduction of microfluidic devices (Stanley et al., 2018). In recent years, microfluidic devices have been employed to image root hairs to enhance live cell imaging (Dupouy et al., 2025; Singh et al., 2021). These recent studies not only made progress in understanding the nuclear morphology during the polarized movement (Singh et al., 2021), but also confirmed the root hair growth dynamics and nuclear movement speed (Dolan et al., 1994; Ketelaar et al., 2002; Singh et al., 2021). They highlighted that the nucleus maintains a mean distance, 77 ± 15 µm, from the growing root hair tip and follows a universal cellular scale for nuclear positioning (Fig. S1). Furthermore, altered agar media has been demonstrated to reduce the root hair growth and nuclear dynamics (Pereira et al., 2024). The calls into question whether the nucleus adheres to the universal scale during varying root hair growth conditions. To determine the position of the nucleus within the root hair, we employed a quantitative approach that measures the developmental stage of the root hair and the intracellular localization of the nucleus. Using live cell imaging, we quantified root hair length (RHL), distance from the base to the nucleus (B2N), and from the nucleus to the root hair tip (N2T) (Fig. 1B). Furthermore, we calculated the B2N/RHL and N2T/RHL ratios to establish the nuclear position relative to root hair development (Fig. 1B). Finally, we also considered B2N/N2T to decipher the relative position of nucleus from both ends of the root hair (Fig. 1B). To understand the spatial relationship between nuclear positioning and root hair growth, we formulated three hypotheses that nuclear positioning may follow within the root hair: (1) no scale, (2) a universal scale, and (3) an internal scale (Fig. S1). The no-scale model occurs when asymmetric cell division happens, the root hair elongates, and the nucleus moves to the end of the root hair, touching the tip (Fig. S1A). The nucleus follows the universal scale when it maintains a constant distance, regardless of the length of the root hair (Fig. S1B). Finally, the internal scale is where the distance from the nuclei to the tip changes depending on the length of the root hair (Fig. S1C).

We manipulated root hair development using hormones, toxic metals, nutrient stress, blocking energy metabolism, and cell wall modification drugs. Our study revealed that nuclear position maintains the internal scale, an adjustable position relative to root hair development. This finding is fundamentally important for basic cell biology applicable to eukaryotic organisms.

## Results

### Nuclear positioning relative to polarized root hair cell development

We set up a quantitative live cell imaging approach to measure the distinct root hair developmental stage and the nuclear position simultaneously (Fig. 1). For this purpose, we utilized the GFP-WIP1 (outer nuclear envelope proteins of Arabidopsis) to visualize the nucleus and counter stained with propidium iodide (PI) to ensure the root hair cells are alive during the treatment, sample preparation, and imaging (Fig. 1B). Seedlings grown on agar media were directly put onto the glass slide, mounted with PI solution, and imaged within 2 minutes. Previous studies demonstrated that the root hair tip growth speed varies between 1 and 2.5 µm/min (Dolan et al., 1994; Singh et al., 2021) and the average nuclear speed is between 0.5 to 1 µm/min (Pereira et al., 2024). Performing live cell imaging within a few minutes ensures the root tip and nucleus are in an endogenous state.

Using the live cell imaging, we measured root hair length (RHL) (Fig. 1D), distance from the base to the nucleus (B2N) (Fig. 1E), and from the nucleus to the root hair tip (N2T) (Fig. 1F). The developing root hair length (RHL) varies between 100 to 800 µm (Fig. 1D). Within these range of developing root hairs, the nuclear position is variable and almost rarely was observed to be touching the tip of the root hair (Fig. 1B). Each of these parameters indicates the root hair length status (RHL) or nuclear positioning (B2N and N2T) (Fig. 1D-F). However, none of these parameters capture the root hair development and the cellular aspect of the nuclear position simultaneously. To circumvent this, we also considered several informative ratios, including B2N/RHL (Fig. 1G), N2T/RHL (Fig. 1H) and B2N/N2T (Fig. 1I). Among these calculated ratios, B2N/RHL and N2T/RHL highlight the nuclear position relative to the root hair development. These parameters help us establish a quantitative aspect to study the nuclear positioning relative to root hair development under control and altered growth environments.

### Elongated root hairs follow the internal scale

To manipulate the root hair length and study the nuclear position scaling, we used a chemical-cell biology approach. According to previous studies, root hairs treated with Indole-3-acetic acid (IAA) are found to have increased root hair length (Rahman et al., 2002). However, it is only after cell type differentiation that auxin functions to promote root hair growth (Masucci and Schiefelbein, 1996). We treated 4-day-old Arabidopsis seedlings with 30nM IAA and found that the 2-day incubation doubles the root hair length (Figure 2A-C). Consequently, the distances from the base to nucleus (B2N) (Fig. S2A) and nucleus to tip (N2T) (Fig. 2D) are altered for the IAA-treated longer root hairs. Interestingly, the B2N/RHL (Fig. S2B), N2T/RHL (Fig. S2E), and B2N/N2T (Fig. S2C) ratios for the control and IAA treatment have similar data distribution. This suggests that while nuclear positioning in IAA-treated roots appears different from the control, the proportional location of the nucleus within the root remains unchanged when examining the ratios. Altogether, it indicates that the nucleus maintains an internal cellular scaling (Fig. S1C).

**Figure 2:**
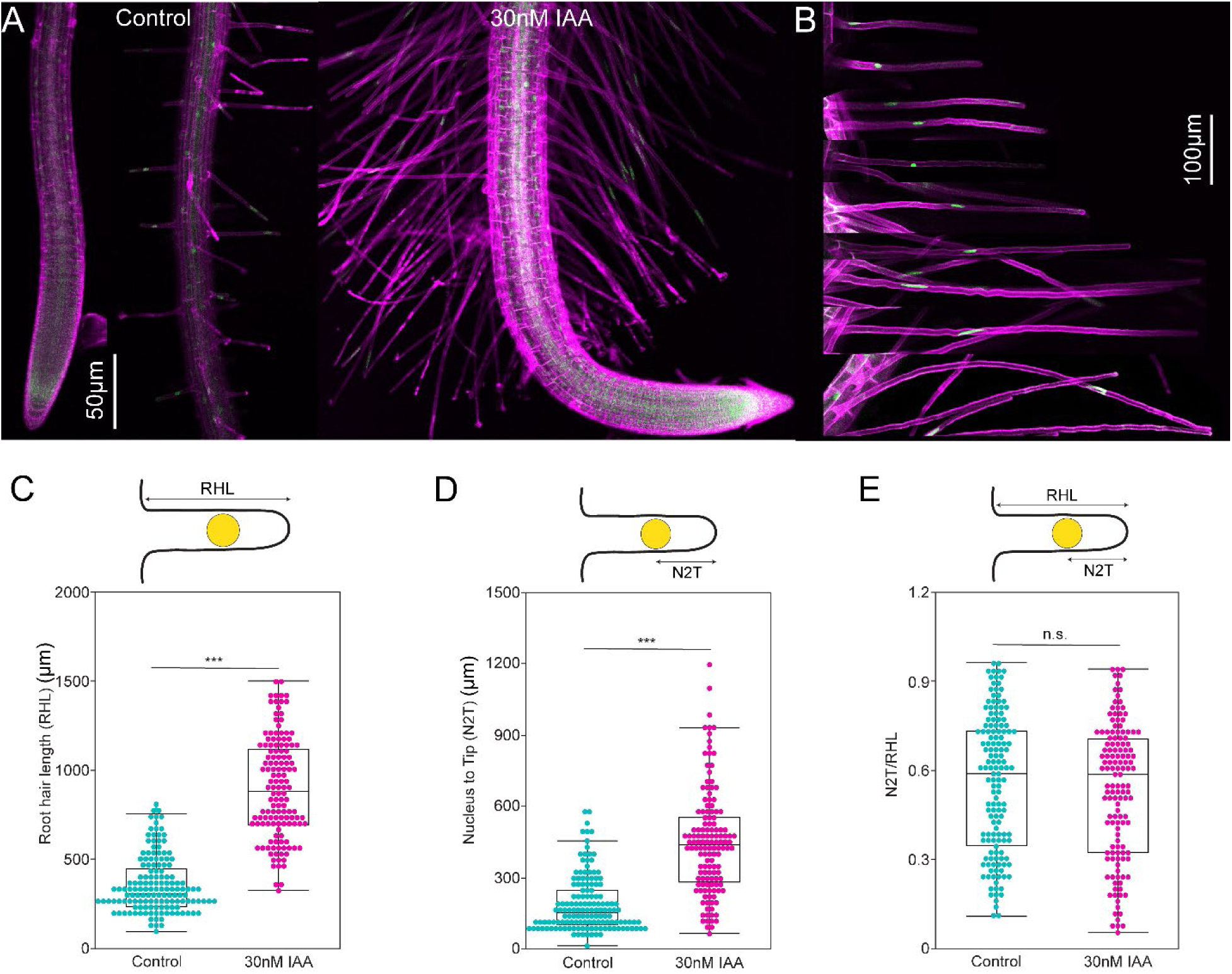
Nuclear positioning in longer root hairs treated with 30nM Indole-3-acetic acid (IAA). (A) Representative 10x images illustrating the phenotypic response of *A. thaliana* to the control treatment and 30nM IAA. Scale = 50 µm. (B) Live cell imaging for root hairs treated with 30nM IAA at varying lengths and nuclear positions. Scale = 100 µm. (C-E) Quantified root hair length (RHL) in control (cyan) versus 30nM IAA-treated root hairs (magenta) (C), measurement of nucleus to tip (N2T) length in control (cyan) and 30nM IAA-treated root hairs (D), and N2T/RHL ratios quantified for control and 30nM IAA-treated root hairs (E). (C-E) The quantification area is shown at the top of each box plot. Each cyan dot represents a root hair that was a control root hair, and each magenta dot represents a root hair treated with 30nM IAA that was measured. Box Plots show median values (center line), 25th to 75th interquartile range (box), and 1.5*interquartile range (whiskers). Student’s t-test was performed for Figure 2 (C-E). Asterisks represent the statistical significance between the means for each treatment. *P < 0.05, **P < 0.01, ***P < 0.001, and n.s. = not significant.

### Nuclear position requires minimum root hair growth for maintaining internal scale

In contrast to the IAA-treated longer root hair, we attempted to create an experimental setup where the root hair length is shorter. Arsenite (As), considered as a toxic metal, is widely distributed, occurring in groundwater, soil, sediment and the atmosphere (Ashraf et al., 2020; Hazelwood et al., 2025a; Smedley and Kinniburgh, 2002). Previous studies demonstrated that 10µM of As caused the primary root length of Arabidopsis to decrease (Ashraf et al., 2020; Bahmani et al., 2016; Hazelwood et al., 2025a), while the root hairs increased in length and density (Bahmani et al., 2016). These studies were performed either by plating the seeds on arsenite-containing media or by transferring seedlings to arsenite-containing plates. The root hair density is usually a product of shorter epidermal cell length. The elongation of root hair, required for water and nutrient uptake, is often an adaptive mechanism when the primary root growth is inhibited under stress conditions. For example, the root hair cell elongates under cold stress, while the primary root growth is inhibited (Urzúa Lehuedé et al., 2025; Zhou et al., 2024).

We transferred 4-days-old seedling to the 10µM As-containing plate and incubated them for 2 days. Unlike findings reported in earlier literature, we found root hairs treated with 10µM As were approximately half the length of those in the control group (Fig. 3A-B). These As-treated shorter root hairs provided the opportunity to study the nuclear position when the polarized growth is inhibited. In As-treated root hair cells, the B2N and N2T distance change as well (Fig. S3A, 3C). Despite differences in nuclear positioning between control and As-treated root hairs, the B2N/RHL (Fig. S3B), N2T/RHL (Fig. 3D), and B2N/N2T (Fig. S3C) reveal a similar proportional relationship. Interestingly, IAA-treated longer root hair (Fig. 2) and As-treated shorter root hair (Fig. 3A-D) most likely follow an internal scale (Fig. S1C), which is adjustable at the cellular level depending on the root hair elongation.

**Figure 3:**
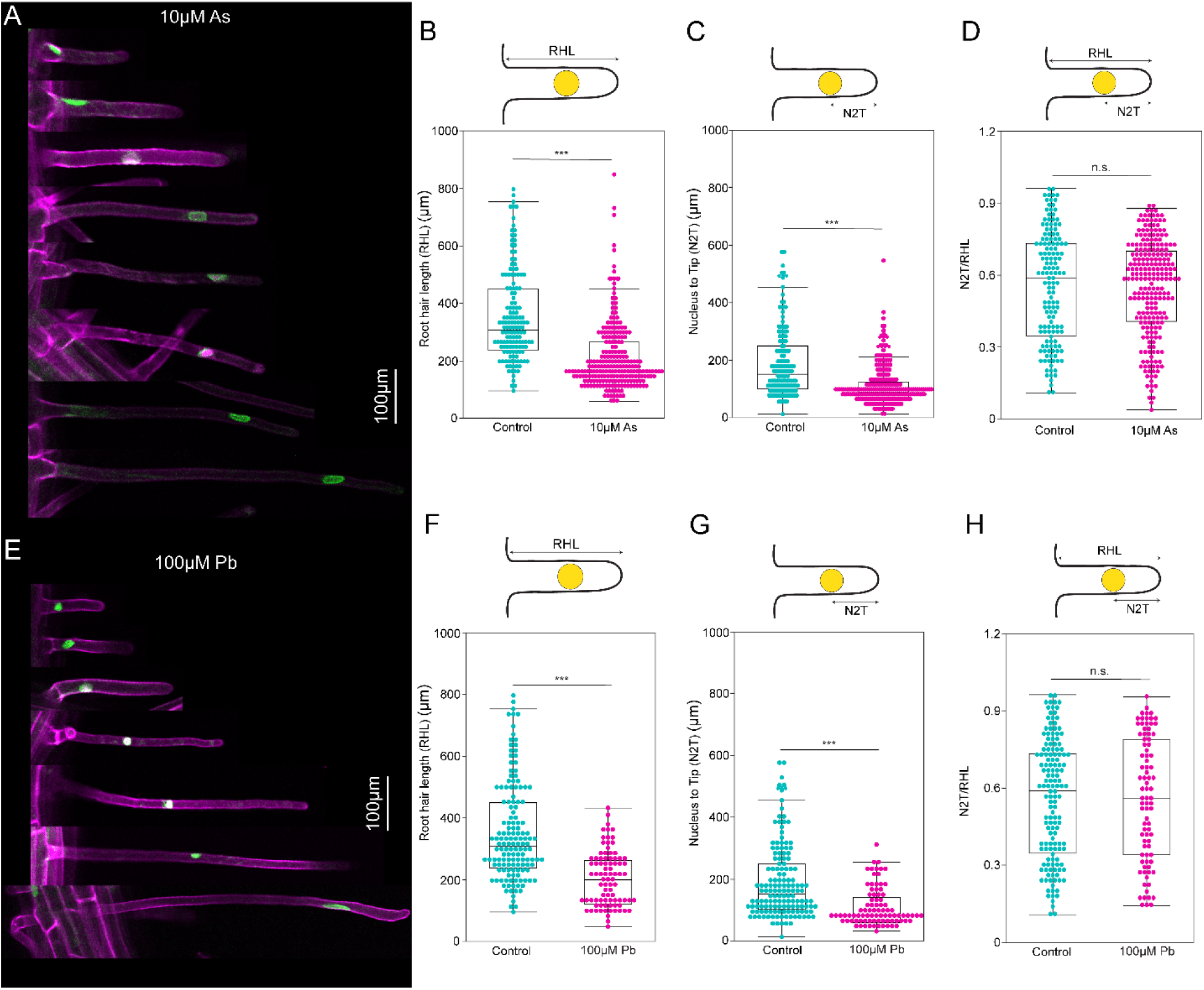
Nuclear positioning in root hairs treated with 10µM arsentie (As) and 100µM lead (Pb). (A) Live cell imaging of representative root hair images treated with 10µM As. Scale = 100 µm. (B-D) Quantification of the root hair length (RHL) (B), nucleus-to-tip distance (N2T) (C), and the N2T/RHL ratio (D) in control versus 10µM As-treated root hairs. (E) Live cell imaging of representative root hair images treated with 100µM Pb. Scale = 100 µm. (F-H) Quantification of RHL (F), N2T (G), and N2T/RHL ratio (H) in control versus 100µM Pb-treated root hairs. (B-H) Quantification areas are indicated at the top of each box plot. Cyan dots represent individual control root hairs, while magenta dots represent root hairs treated with either 10µM As (B-D) or 100µM Pb (F-H). Box Plots show median values (center line), 25th to 75th interquartile range (box), and 1.5*interquartile range (whiskers). Student’s t-test was performed for Figure 3 (B-H). Asterisks represent the statistical significance between the means for each treatment. *P < 0.05, **P < 0.01, ***P < 0.001, and n.s. = not significant.

### The nucleus follows the internal scale while the root hairs grow from a twisted root

The root hair originates from the epidermal cell file, outermost cell layer, of the primary root. During the soil exploration, the primary root experiences gravitropism, waving, curling, twisting, and bending (Porat et al., 2024). Among these root navigation strategies, root twisting is caused by the twisted growth of the epidermal cell layer (Thitamadee et al., 2002). The recent study from our lab suggests that root twisting happens in the presence of toxic metal, lead (Pb) (Hazelwood et al., 2025a). We utilized this knowledge to create shorter root hairs originating from the twisted epidermal cell layer.

We used 4-day-old seedlings to transfer to the 100µM Pb-containing plate and incubated for 2 days. We observed both the twisted root and shorter root hair (Fig. 3E). We quantified the shorter root hair coming out of the twisted part of the root. As a result, the RHL (Fig. 3F), B2N (Fig. S3A), and N2T (Fig. 3G) distances changed for the root hairs treated with 100µM Pb as well. When we converted the parameters into ratios, such as B2N/RHL (Fig. S3B), N2T/RHL (Fig. S3H), and B2N/N2T (Fig. S3C), we unexpectedly found that the data distribution remains the same between the control and 100µM Pb-treated root hairs originating from the twisted primary root. Overall, this result suggests that nuclei maintain the internal scale in mechanically stressed shorter root hair (Fig. S1C).

### Nutrient stress and energy metabolism inhibition-induced shorter root hairs maintain the nuclear internal scale

Root hairs play a crucial structural role in nutrient uptake by increasing the root’s surface area for absorption (Shibata et al., 2022). Arabidopsis is typically grown on 1/2xMS or 1xMS media; however, Keiko Sugimoto’s lab recently demonstrated that increasing the nutrient concentration to 2xMS results in reduced root length (Shibata et al., 2022). This response is thought to help prevent nutrient overloading (Shibata et al., 2022). When we manipulated root hairs with 2xMS, we observed a reduction in their overall length compared to the control (Fig. 4A). As a result of this shortened root hair length, the absolute measurements of B2N (Fig. S4A) and N2T (Fig. 4C) also changed. However, consistent with the other treatments examined, the ratios of B2N/RHL (Fig. S4B), N2T/RHL (Fig. 4D), and B2N/N2T (Fig. S4C) maintained a similar distribution and central tendency. This consistency suggests that nuclei in the 2xMS-treated root hairs are positioned according to an internal scale (Fig. S1C).

**Figure 4:**
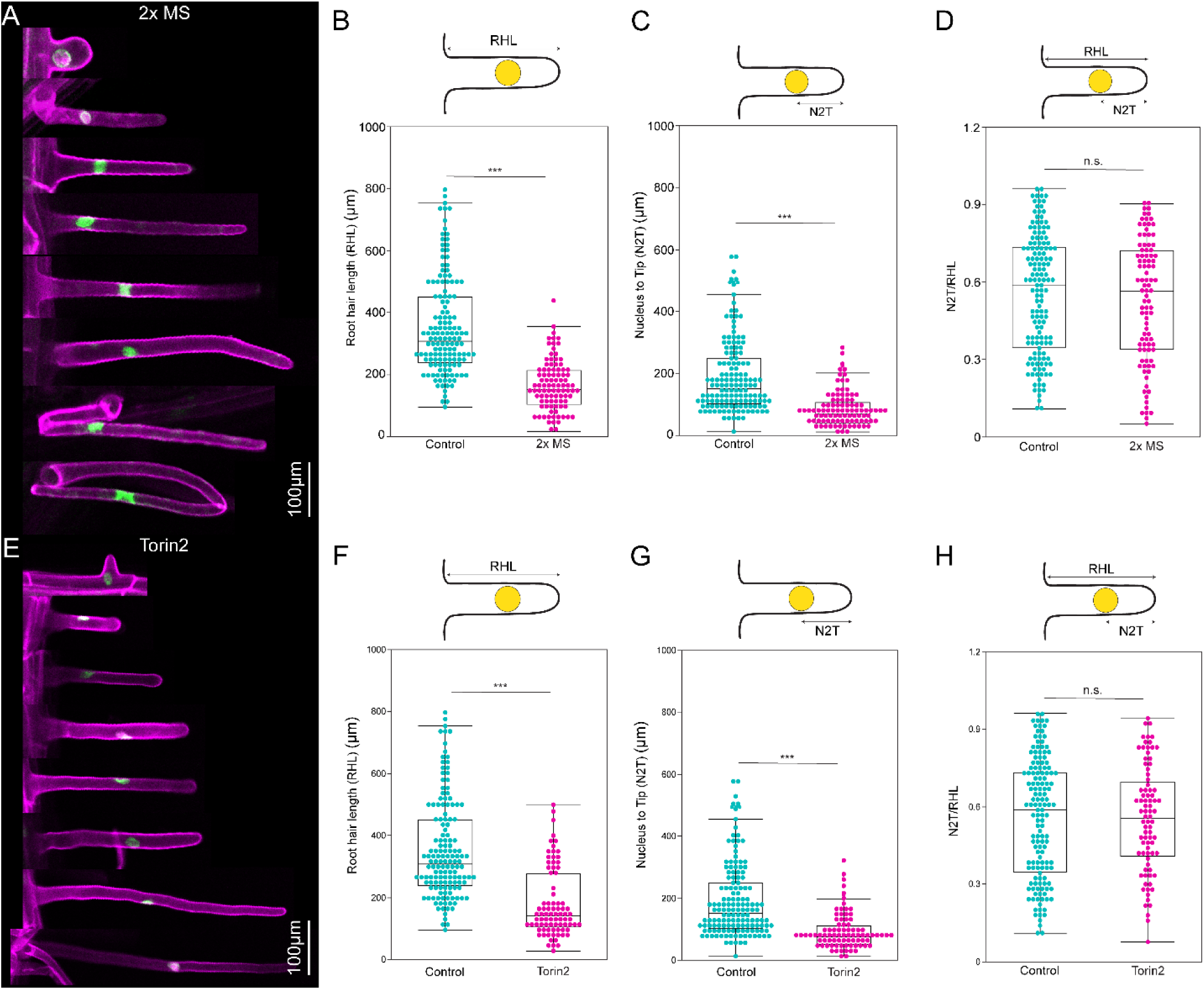
Nuclear positioning in root hairs under 2x MS and Torin2. (A) Live cell imaging of representative root hair images following 2x MS treatment. Scale = 100 µm. (B-D) Comparative analysis of root hair length (RHL) (B), nucleus-to-tip distance (N2T) (C), and the N2T/RHL ratio (D) between untreated controls and 2x MS-treated root hairs. (E) Live cell imaging of representative root hairs after Torin2 exposure. Scale = 100 µm. (F-H) Quantification of RHL (F), N2T (G), and N2T/RHL ratio (H) in control versus Torin2-treated root hairs. (B-H) Each box plot indicates the measurement region at the top. Individual data points are shown in cyan (control) or magenta (2x MS, B-D; Torin2, F-H). Box Plots show median values (center line), 25th to 75th interquartile range (box), and 1.5*interquartile range (whiskers). Student’s t-test was performed for Figure 4 (B-H). Asterisks represent the statistical significance between the means for each treatment. *P < 0.05, **P < 0.01, ***P < 0.001, and n.s. = not significant.

In contrast to excessive nutrients, we looked for an alternative approach to manipulate energy metabolism and create altered root hair development. The TARGET OF RAPAMYCIN (TOR) signaling pathway plays a major role in regulating plant growth and energy metabolism. TOR integrates various internal and external cues such as hormonal, nutrient, light, stress signaling to tailor the plant’s physiological responses according to its overall condition and energy status. It modulates metabolism in response to these factors, thereby optimizing growth and development (Retzer and Weckwerth, 2021). Notably, enhanced TOR activity has been positively correlated with increased root hair growth (Retzer and Weckwerth, 2021). In our study, we used Torin2, a specific inhibitor of TOR kinase, which disrupts the typical functions of the TOR pathway (Montané and Menand, 2013). Previous studies demonstrated that the Torin2 treatment suppresses root hair elongation (Montané and Menand, 2013). We found that treating the root hairs with Torin2 caused the total root hair length to decrease (Fig. 4E-F). Consequently, the B2N (Fig. S6A) and N2T (Fig. 4G) values changed as well. However, when we examined the B2N/RHL (Fig. S6B), N2T/RHL (Fig. 4H), and B2N/N2T (Fig. S6C) ratios, there was a similar data distribution when we compared the Torin2-treated root hairs to the control. This demonstrates that the nuclei in these root hairs follow an internal scale (Fig. S1C) after manipulating the root hair length either with excessive nutrients or blocking a central energy metabolism pathway.

### Isoxaben alters root hair growth, but the nuclear internal scale remains the same

*A. thaliana* cells possess a rigid cell wall outside the plasma membrane, which provides structural integrity and serves multiple biological functions (Vaahtera et al., 2019). The cell wall is essential for growth, development, and defense against abiotic and biotic stress. Isoxaben, a cellulose biosynthesis inhibitor, disrupts cell wall integrity, which can lead to morphological deviations from the wildtype phenotype (Chaudhary et al., 2021; Hoffmann et al., 2024; Scheible et al., 2001). Additionally, Isoxaben has been shown to inhibit root hair elongation through an AVG-sensitive, ethylene-independent pathway (Tsang et al., 2011). In line with previous findings, we observed that Ioxaben not only suppresses root hair length (RHL) (Fig. 5A-B) but also alters the distances from the base to the nucleus (B2N) (Fig. S7A) and from the nucleus to the tip (N2T) (Fig. 5C). Despite these changes, the ratios such as B2N/RHL (Fig. S7B), N2T/RHL (Fig. 5D), and B2N/N2T (Fig. S7C) remained consistently similar to the control. Alteration of root hair development through cellulose biosynthesis inhibitor Isoxaben maintains the internal scale for nuclear position (Fig. S1C).

**Figure 5:**
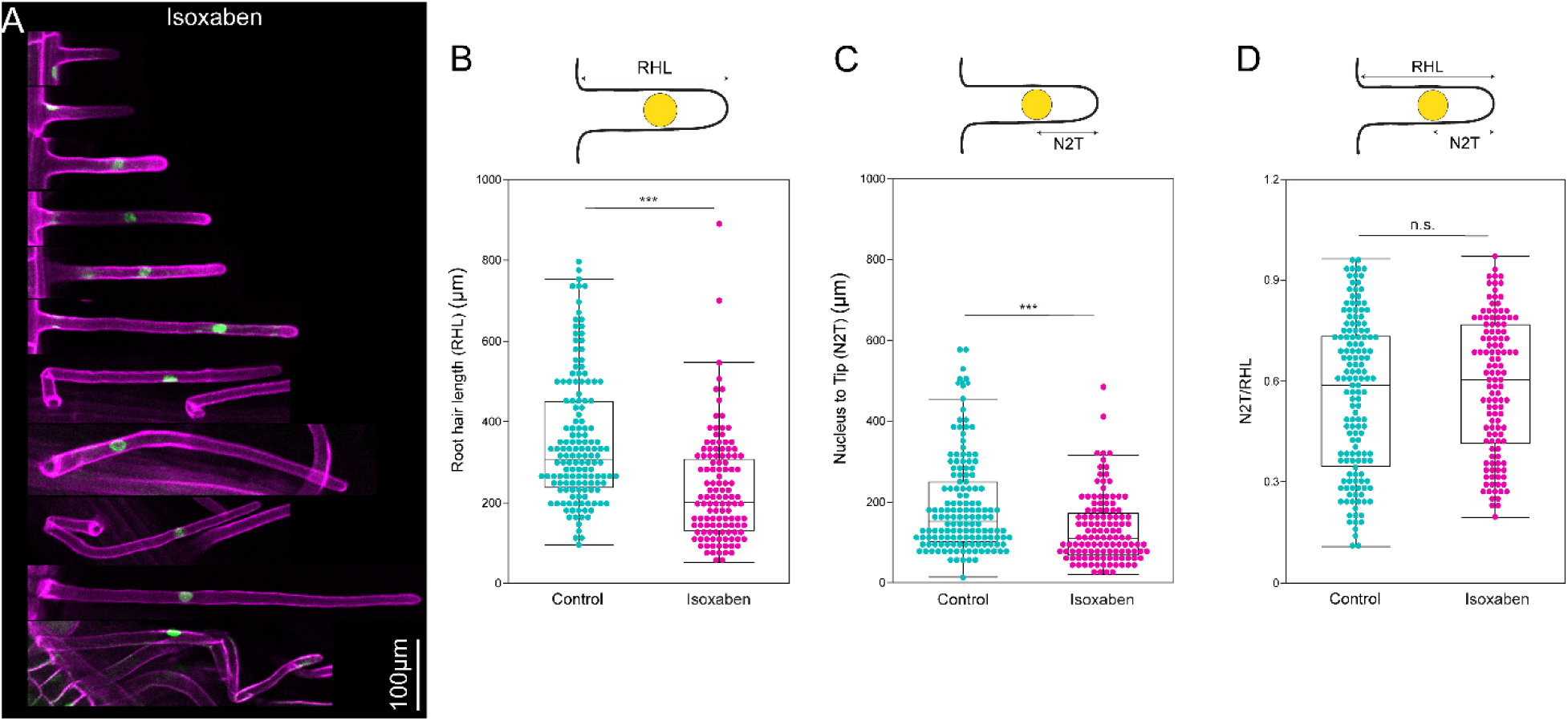
Nuclear positioning in root hairs exposed to isoxaben (Ixb). (A) Representative live-cell image of a root hair following Ixb treatment. Scale = 100 µm. (B-D) Analysis of the root hair length (RHL) (B), nucleus-to-tip distance (N2T) (C), and the N2T/RHL ratio (D) compared untreated controls and Ixb-treated root hairs. (B-D) Cyan dots indicate individual control measurements, whereas magenta dots correspond to Ixb-treated root hairs. Box Plots show median values (center line), 25th to 75th interquartile range (box), and 1.5*interquartile range (whiskers). Student’s t-test was performed for Figure 5 (B-D). Asterisks represent the statistical significance between the means for each treatment. *P < 0.05, **P < 0.01, ***P < 0.001, and n.s. = not significant.

At this point, our focus turned to examining the intriguing scale of nuclear positioning. No prior reports of internal nuclear scaling in polarized cells were reported, so we looked for examples beyond the cellular aspects. Across all treatments, including IAA, As, Pb, 2xMS, Torin2, and Isoxaben, the root hair exhibited variability in length (Fig. 2-5). However, when analyzing the calculated ratios, we observed remarkable consistency: the distribution and convergence points of these ratios (B2N/RHL, N2T/RHL and B2N/N2T) in treated samples closely matched those in the controls (Fig. 2-5). Notably, the B2N/RHL ratio consistently clustered around approximately 0.4 regardless of the treatment conditions. This persistence of a stable ratio under diverse and dynamic conditions was striking, and it prompted us to investigate its biological significance. Upon further exploration, specifically by inverting the N2T/RHL ratio, we discovered that the central tendency of the data converged ∼1.61 (Fig. S8). This is a mathematically and naturally significant number known as the “Golden ratio” (Marples and Williams, 2022). For the first time, this finding revealed the Golden ratio demonstrated at the cellular level.

### Breaking the internal scale in branched root hair cells

To test the hypothesis of whether polarized growth of the root hair is required for maintaining the internal nuclear positioning scale, we created a branched root hair. The branched root hairs simulate trichomes, where the nucleus is observed at the base of a branched unicellular cell (Hülskamp and Mathur, 2001). A previous study suggested that 100nM epibrassinolide (eBL) alters the epidermal cell patterning and the root hair (Cheng et al., 2014). We observed split root hairs after treating Arabidopsis seedlings with 100nM eBL for 2 days (Fig. 6A). Our quantification data demonstrated that more than 50% of root hairs show a branched phenotype (Fig. 6B). Next, we observed the nuclear position within split root hairs and found nuclear position in the base (∼20%) or shorter branch (∼35%) or longer branch (∼45%) of the cell (Fig. 6C). This observation and quantitative data indicate that it is the unidirectional polarized growth of root hairs – rather than their length – that is required to maintain the internal scale of nuclear positioning.

**Figure 6:**
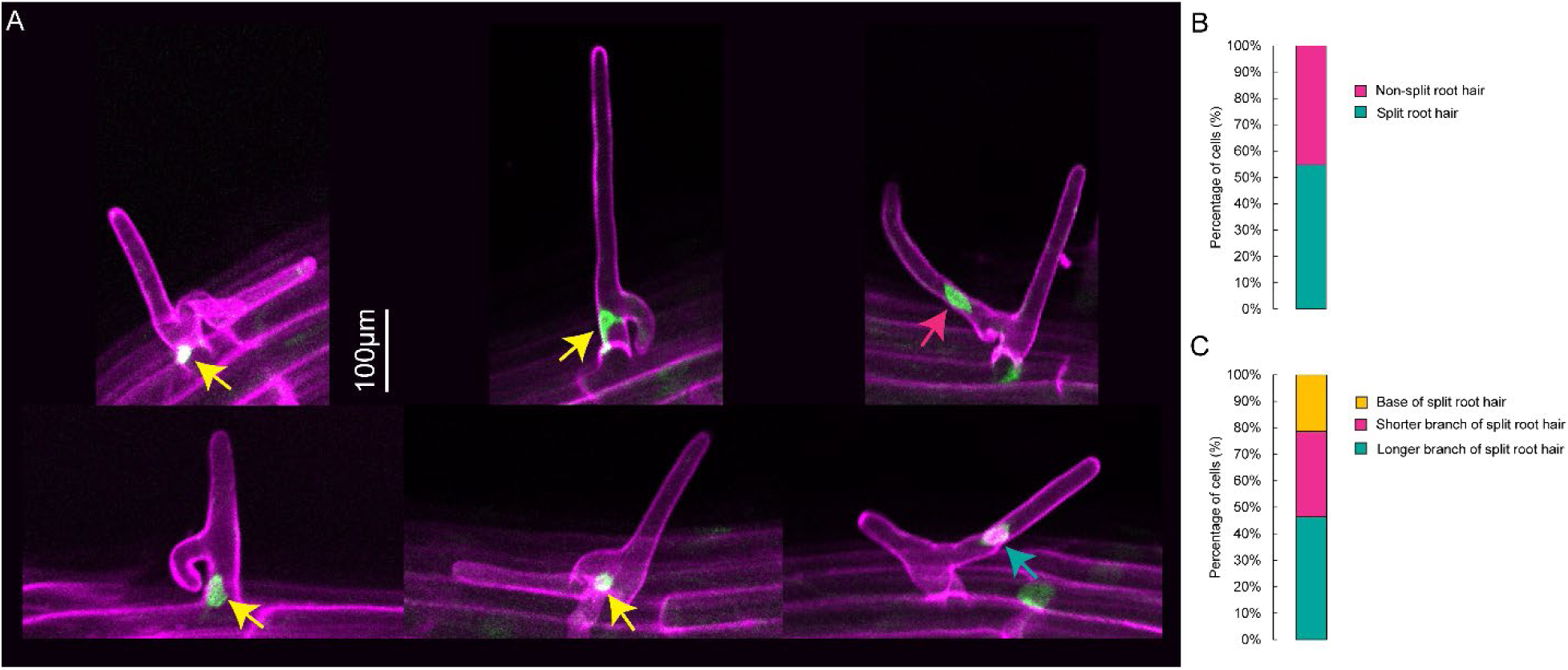
Nuclear positioning in root hairs treated with 24-epibrassinolide (eBL). (A) Live cell imaging of representative root hair images treated with eBL. Scale = 100 µm. (B) Proportion of eBL-treated root hair cells that developed a split phenotype (cyan) versus those that remain unsplit (magenta). (D) Percentage of the distribution of nuclear positioning within split root hairs: nuclei located at the base of the split (yellow), within the shorter branch (magenta), or the longer branch (cyan).

## Discussion

Our study discovered that the nucleus maintains an internal spatial cellular scale during polarized root hair growth, and the scale remains intact even during the altered root hair growth (Fig. 7). This finding is fundamentally important for nuclear positioning during polarized growth.

**Figure 7:**
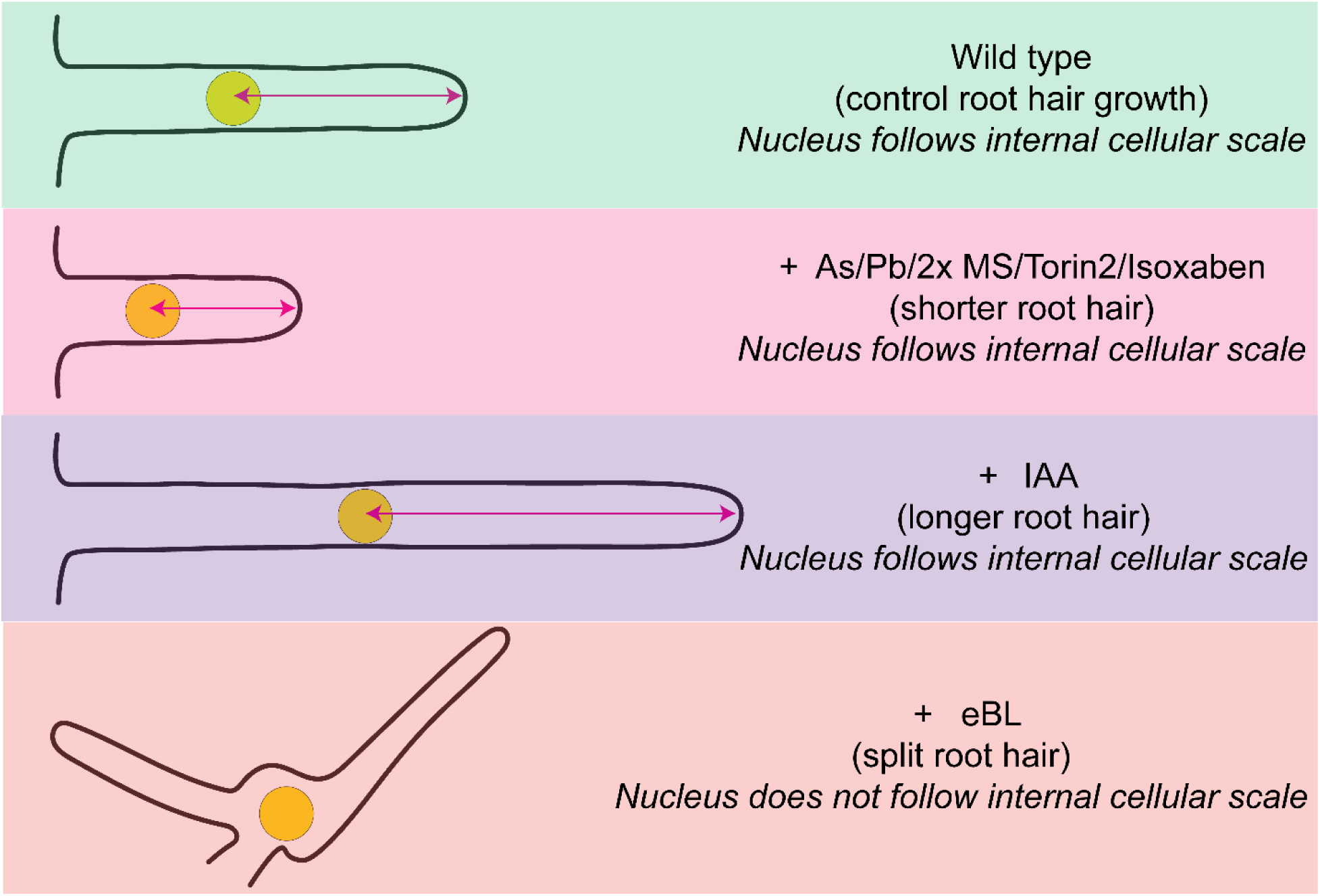
Models of root hair developmental stages under different treatments and their associated nuclear positioning. (A) Model of a control root hair, where the nucleus positions itself according to an internal scale. (B) Model of root hairs treated with As, Pb, 2x MS, Torin2, or isoxaben, which are shorter but still show nuclear positioning that follows an internal cellular scale. (C) Model of IAA-treated root hairs, which are longer and also maintain nuclear positioning along the internal scale. (D) Model of eBL-treated root hairs, which develops a split phenotype in which the nucleus no longer follows an internal scale.

We carefully assessed the previously published articles about nuclear positioning in *A. thaliana* root hair cells from the past three decades. During this process, we have identified that (a) the time length used for time-lapse imaging was not sufficient, and (b) the sample size was lacking to adequately capture the nuclear dynamics during the entire root hair development process. For instance, the root hair growth, which starts with bulge formation and elongates up to ∼1000 µm with an average speed of ∼1.5 µm/min, is a continuous process (Dolan et al., 1994; Ketelaar et al., 2002; Singh et al., 2021). In that case, we need to have a time lapse for ∼11h (total length ∼1000 µm / average speed ∼1.5 µm/min = ∼666 min) to capture the nuclear position during the entire root hair growth from the beginning to the end. The published literature reported the time lapse studies with a maximum of 60 minutes (Brueggeman et al., 2022; Ketelaar et al., 2002; Singh et al., 2021). Within this period of 60 minutes, the root hair grows ∼90 µm and it represents ∼10% of the entire root hair development. To circumvent this issue, we relied on measuring nuclear position from a wide range of developing root hairs, ranging between 100 µm and 800 µm (Fig. 1-6). Additionally, rather than considering the nuclear dynamics or nuclear morphology or tip growth speed individually, we calculated the spatial relationship between nuclear position and root hair development through B2N/RHL and N2T/RHL ratios.

The use of a microfluidic device enhanced the possibility of live cell imaging of root hairs (Dupouy et al., 2025; Pereira et al., 2024; Singh et al., 2021; Stanley et al., 2018). At the same time, the device causes three major issues: (a) it does not replicate the actual root hair growth environment; (b) it is not possible for all of the root hairs to enter the microfluidic channel; and (c) we are not able to study the much longer root hair cells.

The microfluidic device provides an artificial path for the root hair, where the only growth can occur in a straight line. In reality, root hairs are generated from a 3D cylindrical primary root’s epidermal cell layers and grow in all possible directions. In our study, we considered the root hairs growing all possible 3D surfaces of the epidermal cell layers, including even folded and bent root hairs. This attempt ensures to capture the root hair, as well as the nuclear position in a way similar to their native growth conditions.

We believe our study provides a foundational framework for understanding the spatial relationship between nuclear position and polarized root hair cell growth. The identified internal cellular scale will be instrumental for future genetic screening, environmental stress response, and examining crop varieties. Furthermore, the central tendency of the internal scale towards the Golden ratio has laid the groundwork for mathematical modeling and exploring the nuclear position in polarized cells beyond the root hair in *A. thaliana*.

## Methods and materials

### Plant materials

*Arabidopsis thaliana* seeds GFP-WIP1 (ABRC #CS39987) were used for nuclear visualization.

### Plant growth conditions

For Arabidopsis, seeds were surface sterilized with 1mL 70% ethanol for 10 minutes, washed twice with 1mL autoclaved MQ water inside the laminar flow hood, and placed on the surface of ½ Murashige and Skoog (MS) media (BioWorld; Cat.# 30630058) containing 1% sucrose (BioWorld; Cat.# 41900152) and 1% agar (BioWorld; Cat.# 40100072) in square petri dishes (Simport; Cat.# 26-275). Plate openings were sealed with micropore tapes (Amazon; 3M A-1530-0 Micropore Surgical Tape, ½ Inch), covered with aluminum foil, and kept at 4°C for 2 days. After 2 days, plates are placed vertically under constant light conditions at 22°C.

### Chemical treatments

4-days-old Arabidopsis seedlings were transferred to 30nM IAA (BioWorld; Cat.# 30631010-1), 10uM As (Sigma-Aldrich; Cat.# S7400), 100uM Pb (Sigma-Aldrich; Cat.# 228621), 2x MS (BioWorld; Cat.# 30630058), Isoxaben (Sigma-Aldrich; Cat.# 36138), Torin2 (Cayman; Cat.# 14185), 300uM ZnSO4 (Fisher Scientific; Cat.# Z68-500), 24-Epibrassinolide or eBL (Cayman; Cat.# 20337); and incubated for 2 days before imaging.

### Microscopy

GFP and propidium iodide (PI) fluorescence was viewed and captured using a Nikon Eclipse 80i confocal laser-scanning microscope or an Olympus IXplore SpinSR system (Evident, Tokyo, Japan), equipped with an inverted microscope (IX83; Evident, Japan), a CSUW1-Sora spinning disk confocal unit (Yokogawa, Japan) and a Hamamatsu ORCA Fusion BT camera. For the Nikon Eclipse 80i confocal laser-scanning microscope, laser lines 488nm and 543nm laser lines were used for GFP and PI, respectively. For the Olympus IXplore SpinSR system, 488nm and 561nm laser lines were used for GFP and PI, respectively. Samples were observed and images were captured either using 10x or 20x objectives.

### Image analysis

The open-source ImageJ/Fiji software software (<u>imagej.nih.gov/ij/</u>) was used to measure the root hair length (RHL), base to nucleus (B2N), and nucleus to tip (N2T). The distance is measured from the center of the nucleus. The representative images are pseudo-colored, where GFP-WIP1 and PI are highlighted as green and magenta, respectively; using ImageJ/Fiji software.

### Statistical analysis

The raw data for each quantification was imported to statistical software JMP Pro17 for generating graphs and performing statistical tests.

## Acknowledge

The authors thank Abel Rosado for providing access to the Nikon Eclipse 80i confocal laser-scanning microscope, Sean Ritter from Wasteneys’ lab for sharing chemicals, Miki Fujita and EunKyoung Lee of UBC Bioimaging Facility (RRID: SCR_021304) for their kind support.

---

**Figure S1:**
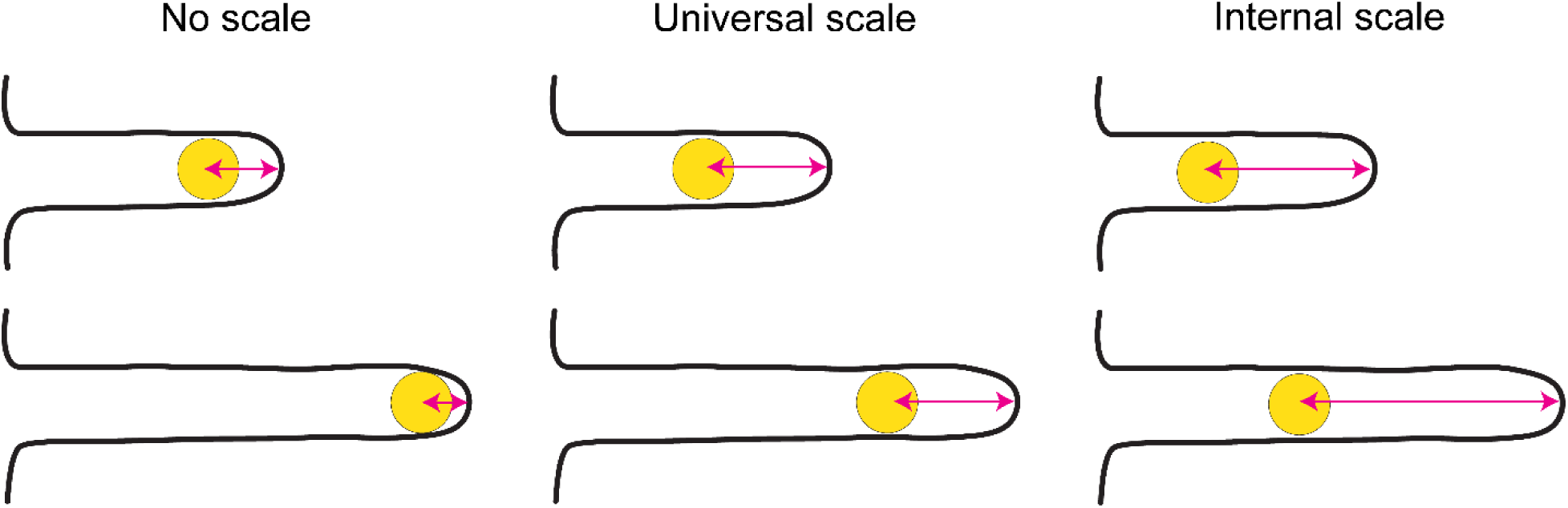
Models of nuclear positioning in root hairs. There are three patterns: (i) no scale, where the nucleus is positioned at the end of the root hair; (ii) universal scale, where the nucleus maintains a fixed distance from the tip regardless of root hair length; and (iii) internal scale, where the nucleus-to-tip distance changes in proportion to root hair length.

**Figure S2:**
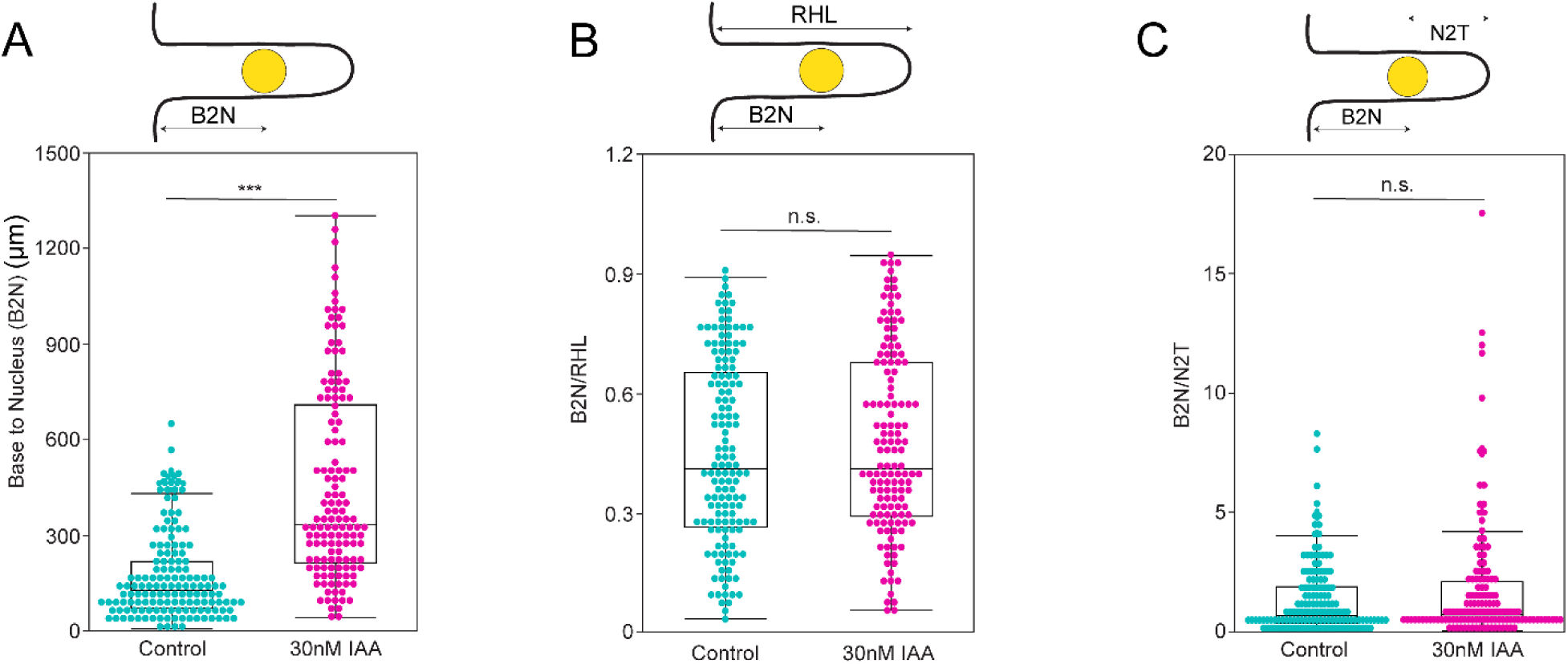
Quantitative measurement of nuclear positioning and root hair in presence of IAA. (A-C) Quantified measurement of the base to the nucleus length (B2N) in control (cyan) versus 30nM IAA-treated root hairs (magenta) (A), B2N/RHL ratios quantified for control and 30nM IAA-treated root hairs (B) and B2N/N2T ratios quantified for control and 30nM IAA-treated root hairs (C). (A-C) The quantification area is shown at the top of each box plot. Each cyan dot represents a root hair that was a control root hair, and each magenta dot represents a root hair treated with 30nM IAA that was measured. Box Plots show median values (center line), 25th to 75th interquartile range (box), and 1.5*interquartile range (whiskers). Student’s t-test was performed for Figure 2 (C-E). Asterisks represent the statistical significance between the means for each treatment. *P < 0.05, **P < 0.01, ***P < 0.001, and n.s. = not significant.

**Figure S3:**
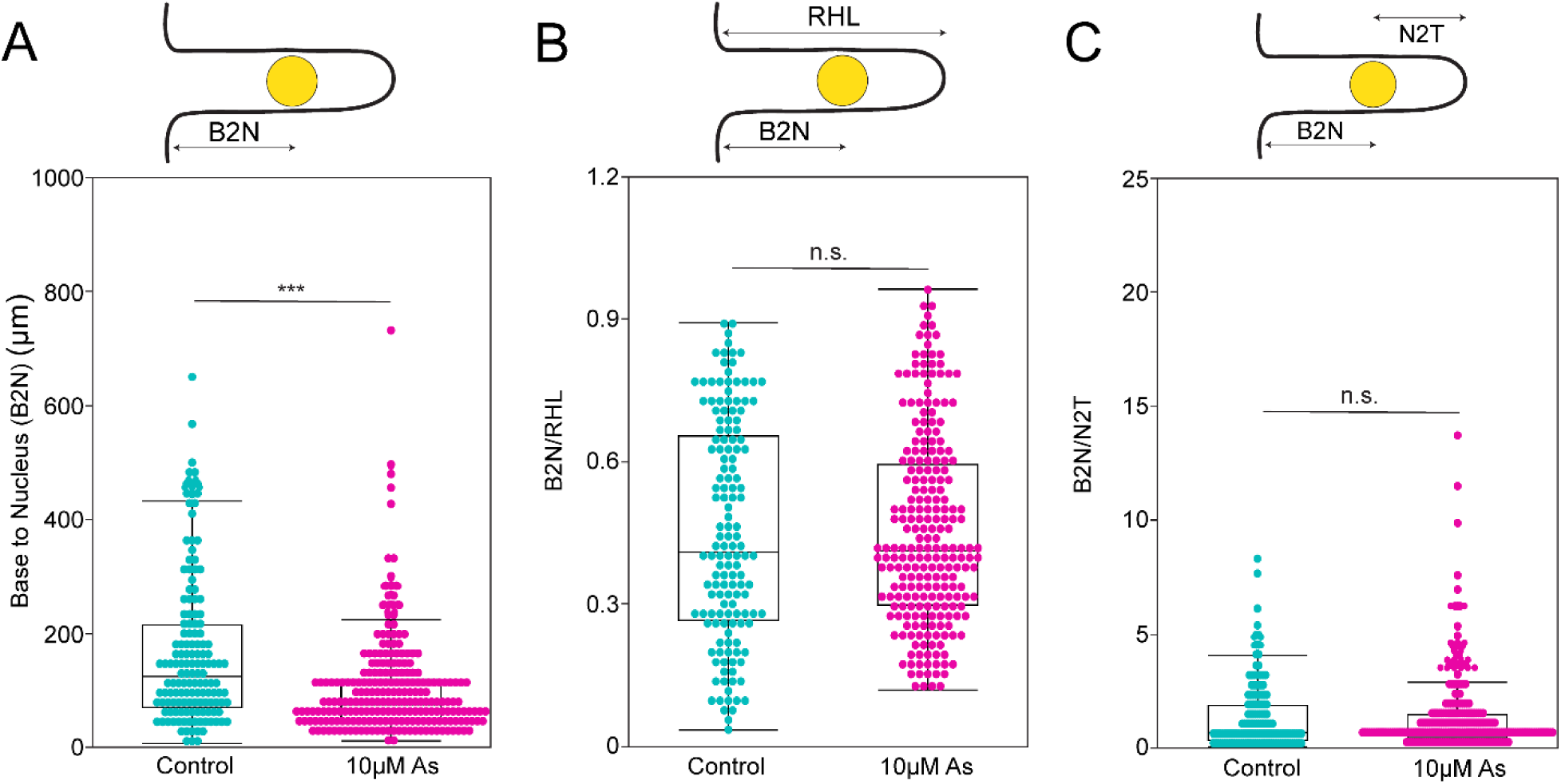
Quantitative measurement of nuclear positioning and root hair in presence of As. (A-C) Measurements of nuclear positioning in control and 10µM arsenite (As) treated root hairs for the base-to-nucleus (B2N) length (A), B2N/RHL ratio (B), and B2N/N2T ratio (C). Quantification areas are indicated at the top of each box plot. Each point represents an individual root hair (cyan = control, magenta = 10 µM As). Box Plots show median values (center line), 25th to 75th interquartile range (box), and 1.5*interquartile range (whiskers). Student’s t-test was performed for Figure 2 (C-E). Asterisks represent the statistical significance between the means for each treatment. *P < 0.05, **P < 0.01, ***P < 0.001, and n.s. = not significant.

**Figure S4:**
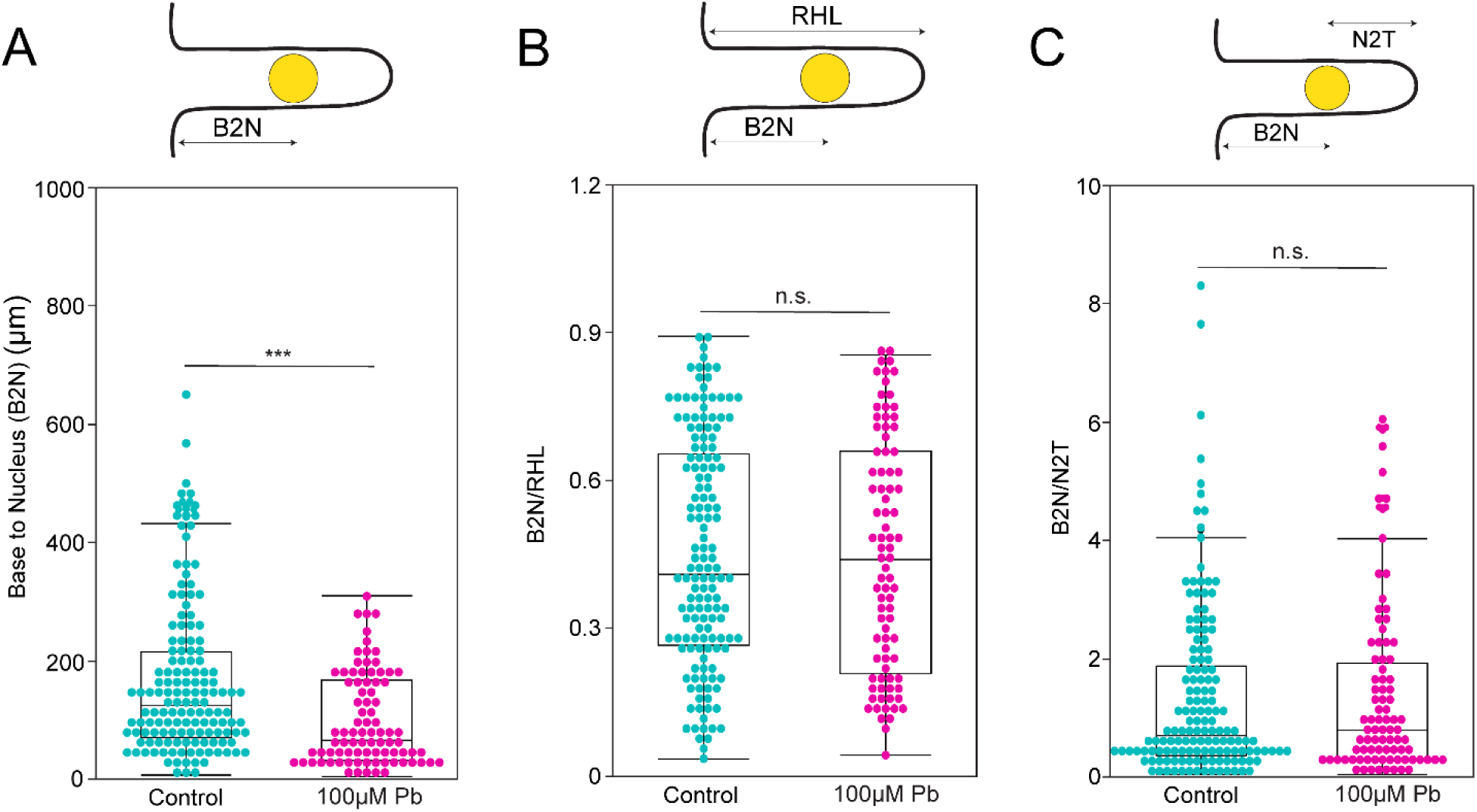
Quantitative measurement of nuclear positioning and root hair in presence of Pb. (A-C) Analysis of nuclear positioning in control versus 100µM lead (Pb) treated root hairs. Shown are (A) base-to-nucleus (B2N) length, (B) B2N/RHL ratio and (C) B2N/N2T ratio for control-treated root hairs and 100µM Pb-treated root hairs. The region quantified is indicated at the top of each box plot. Individual data points correspond to single root hairs (cyan = control, magenta = 100µM Pb). Quantification areas are indicated at the top of each box plot. Each point represents an individual root hair (cyan = control, magenta = 100 µM Pb). Box Plots show median values (center line), 25th to 75th interquartile range (box), and 1.5*interquartile range (whiskers). Student’s t-test was performed for Figure 2 (C-E). Asterisks represent the statistical significance between the means for each treatment. *P < 0.05, **P < 0.01, ***P < 0.001, and n.s. = not significant.

**Figure S5:**
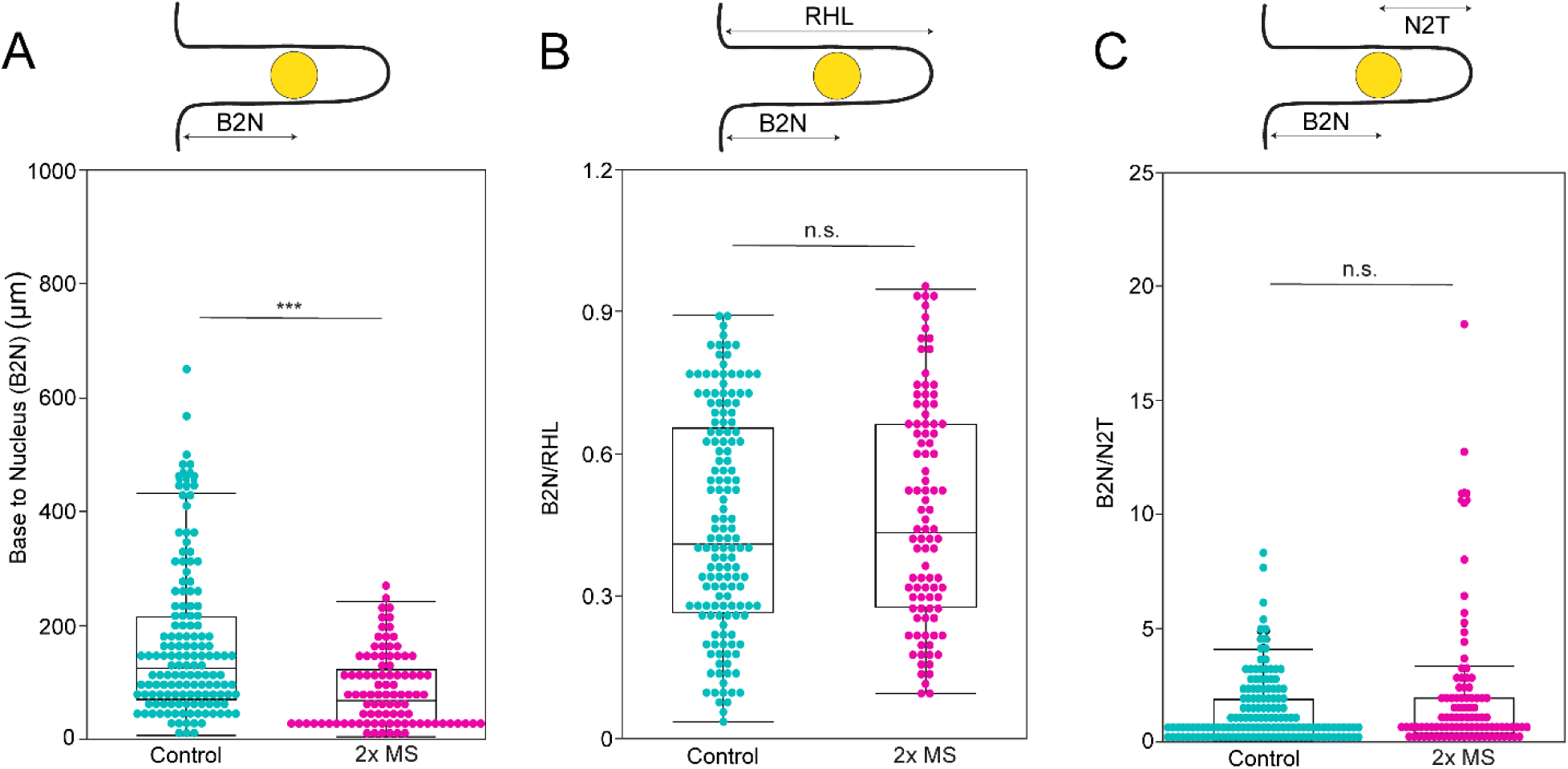
Quantitative measurement of nuclear positioning and root hair in presence of 2xMS. (A-C) Nuclear position in root hairs is subjected to standard conditions or 2x MS treatments. Parameters measured are (A) base-to-nucleus length (B2N), (B) B2N/RHL ratio, and (C) B2N/N2T ratio. The quantified measurement range is indicated above each panel. Each point corresponds to one root hair, with cyan denoting control and magenta representing 2x MS treatment. Box Plots show median values (center line), 25th to 75th interquartile range (box), and 1.5*interquartile range (whiskers). Student’s t-test was performed for Figure 2 (C-E). Asterisks represent the statistical significance between the means for each treatment. *P < 0.05, **P < 0.01, ***P < 0.001, and n.s. = not significant.

**Figure S6:**
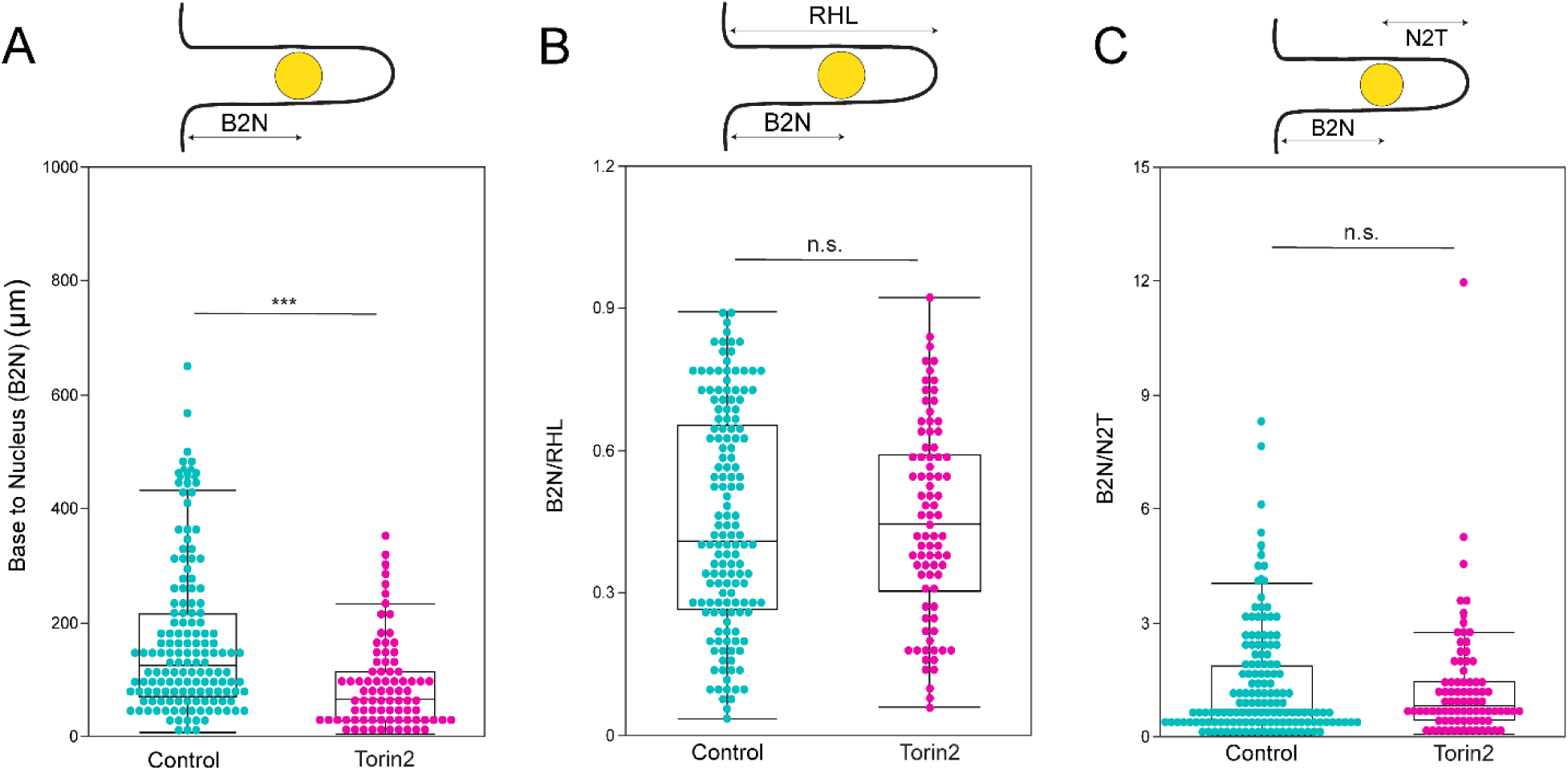
Quantitative measurement of nuclear positioning and root hair in presence of Torin2. (A-C) Evaluation of nuclear positioning parameters in control versus Torin2-treated root hairs. Plotted values include (A) base-to-nucleus length (B2N), (B) B2N/RHL ratio, and (C) B2N/N2T ratio. The specific region quantified is noted above each box plot. Data points represent individual root hairs, where cyan is control-treated root hairs and magenta is Torin2-treated root hairs. Box Plots show median values (center line), 25th to 75th interquartile range (box), and 1.5*interquartile range (whiskers). Student’s t-test was performed for Figure 2 (C-E). Asterisks represent the statistical significance between the means for each treatment. *P < 0.05, **P < 0.01, ***P < 0.001, and n.s. = not significant.

**Figure S7:**
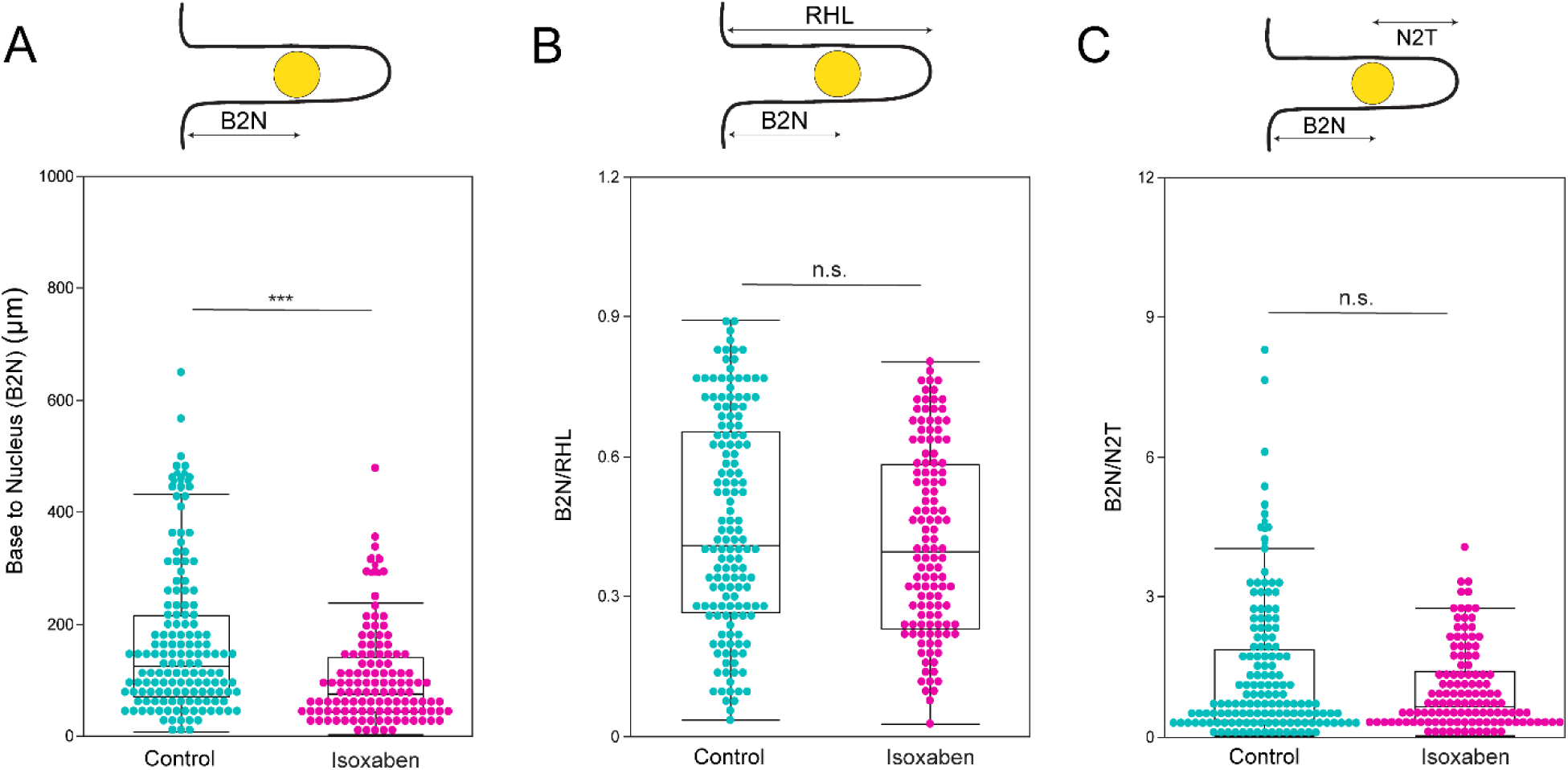
Quantitative measurement of nuclear positioning and root hair in presence of Isoxaben. (A-C) Analyzed root hairs exposed to Isoxaben (Ixb) for changes in nuclear positioning relative to untreated controls. The three parameters measured are (A) base-to-nucleus distance (B2N), (B) the ratio of B2N to overall root hair length (RHL) (B2N/RHL ratio), and (C) the ratio of B2N to nucleus-to-tip distance (N2T) (B2N/N2T ratio). The measured area is indicated above each of the plots. Data points represent individual root hairs, where cyan is control-treated root hairs and magenta is Ixb-treated root hairs. Box Plots show median values (center line), 25th to 75th interquartile range (box), and 1.5*interquartile range (whiskers). Student’s t-test was performed for Figure 2 (C-E). Asterisks represent the statistical significance between the means for each treatment. *P < 0.05, **P < 0.01, ***P < 0.001, and n.s. = not significant.

**Figure S8:**
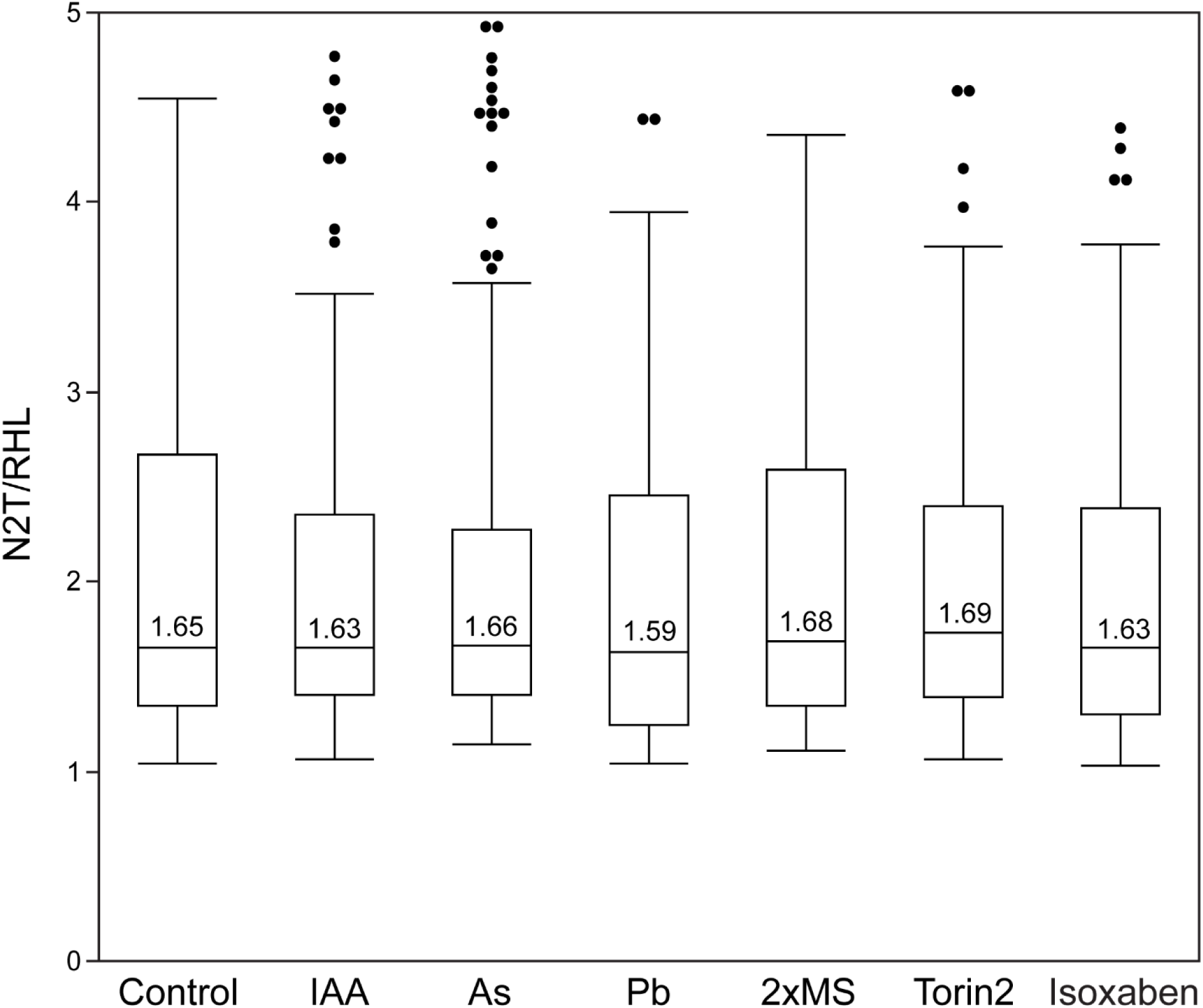
Box plots of nuclear positioning in *A. thaliana* root hairs under control, IAA, As, Pb, 2x MS, Torin2 and Isoxaben treatments. The nucleus-to-tip/total root hair length ratio shows a consistent convergence around 1.61 across conditions.

