## Supplementary figures and images for "The nucleus follows an internal cellular scale during polarized root hair cell development"

### Supplemental FIgure 1.tif

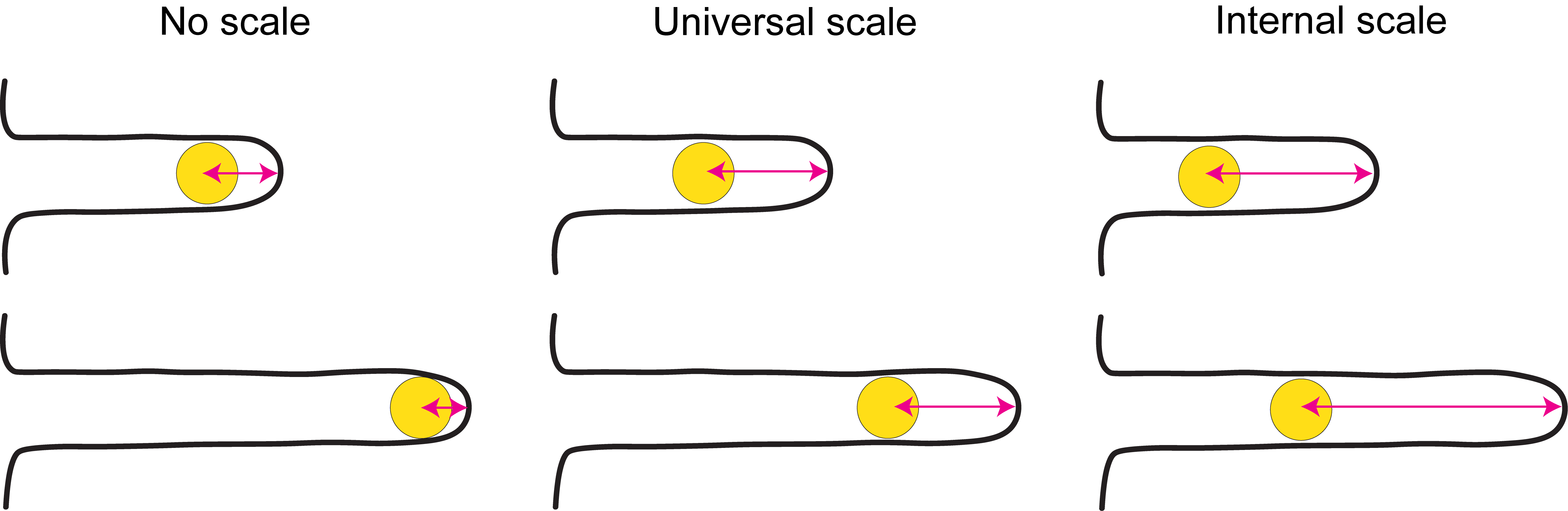

### Supplemental Figure 2.tif

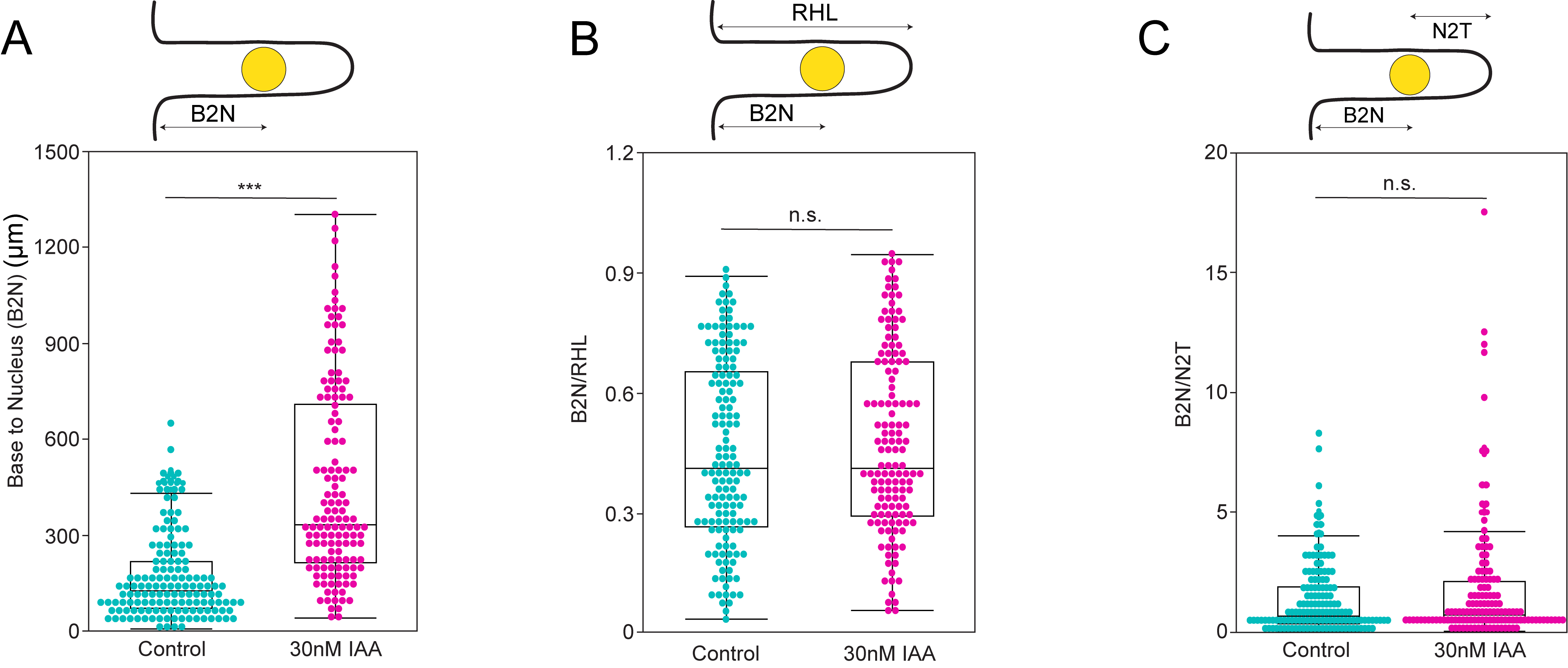

### Supplemental Figure 3.tif

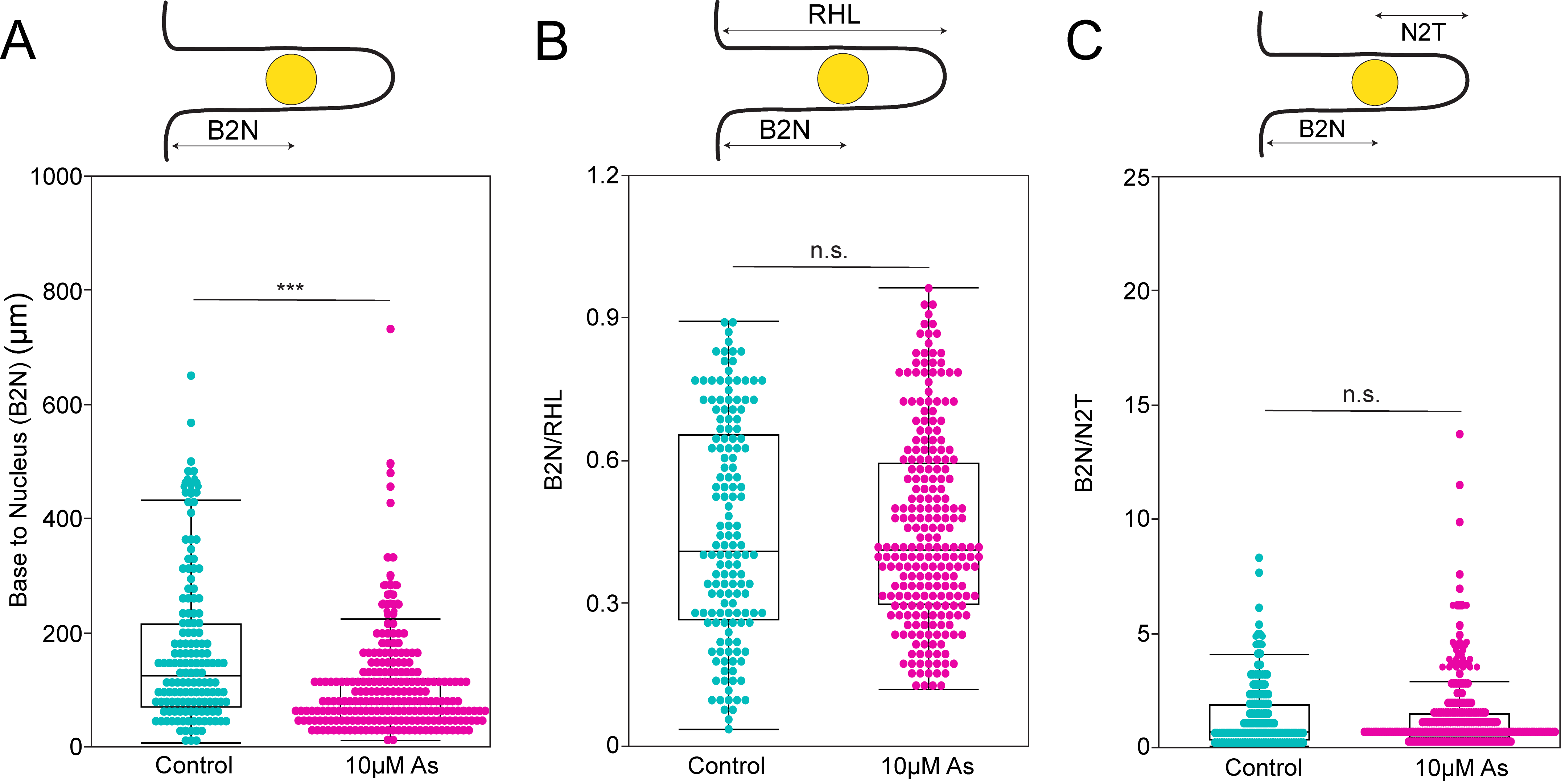

### Supplemental Figure 4.tif

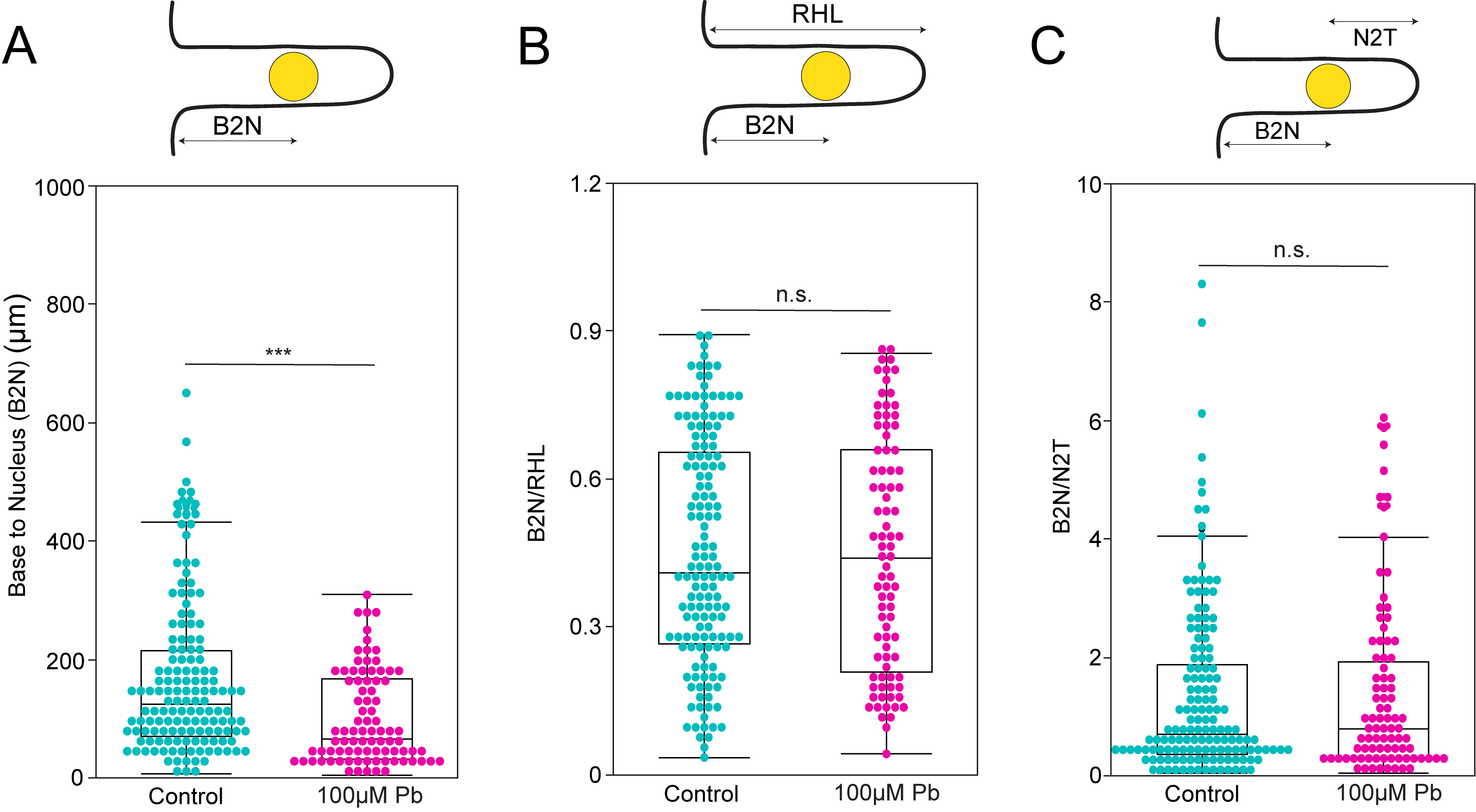

### Supplemental Figure 5.tif

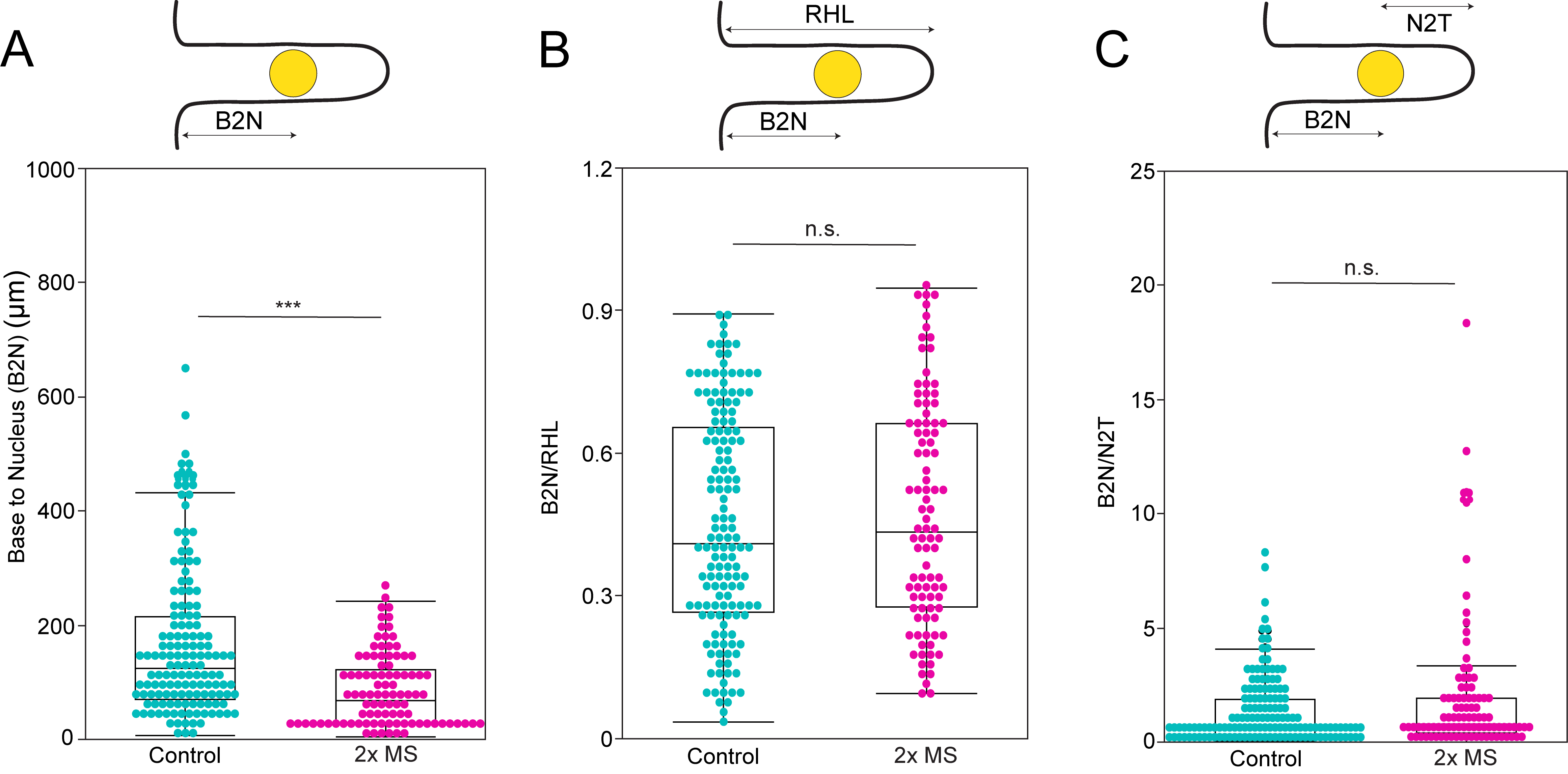

### Supplemental Figure 6.tif

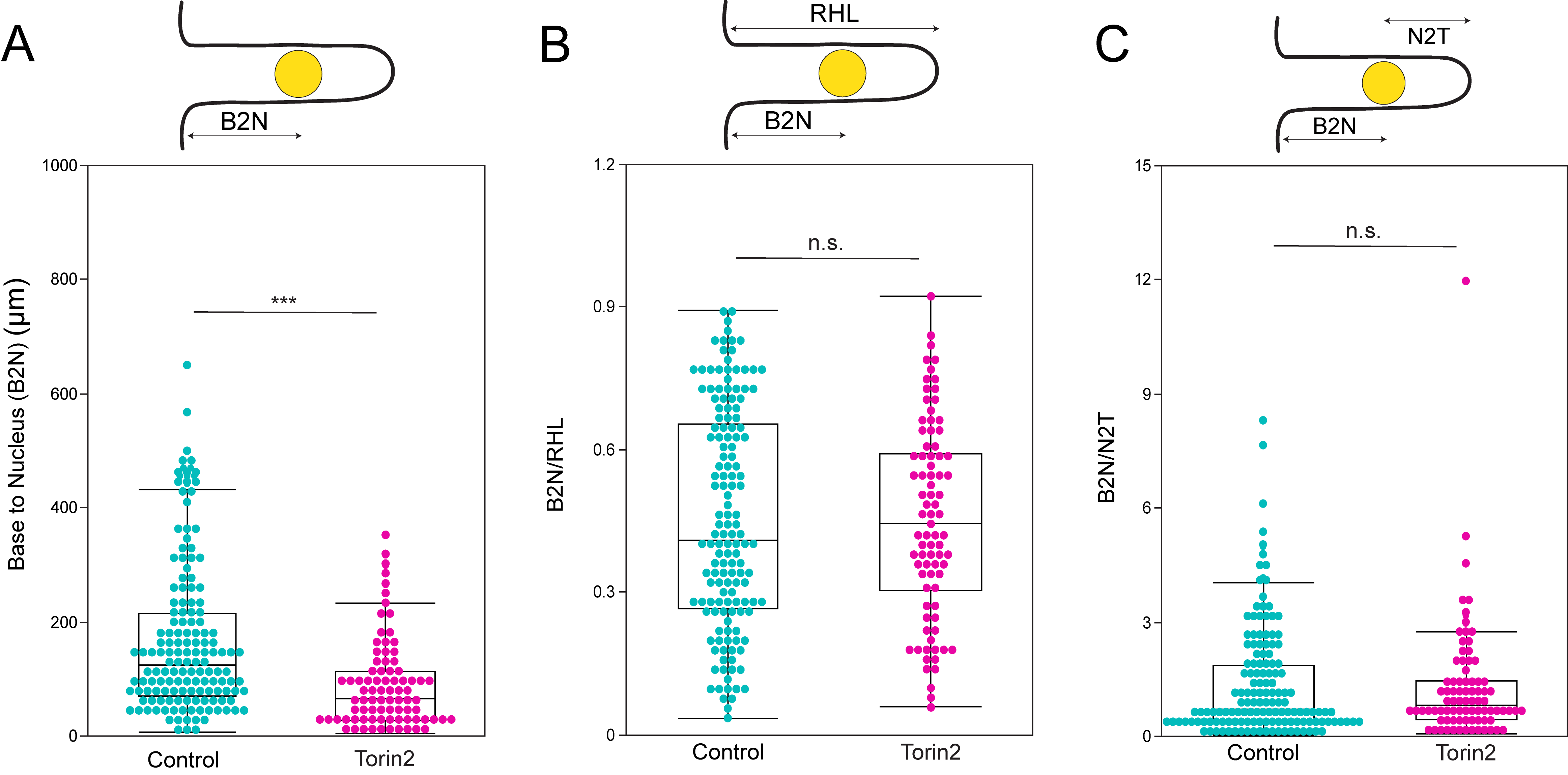

### Supplemental Figure 7.tif

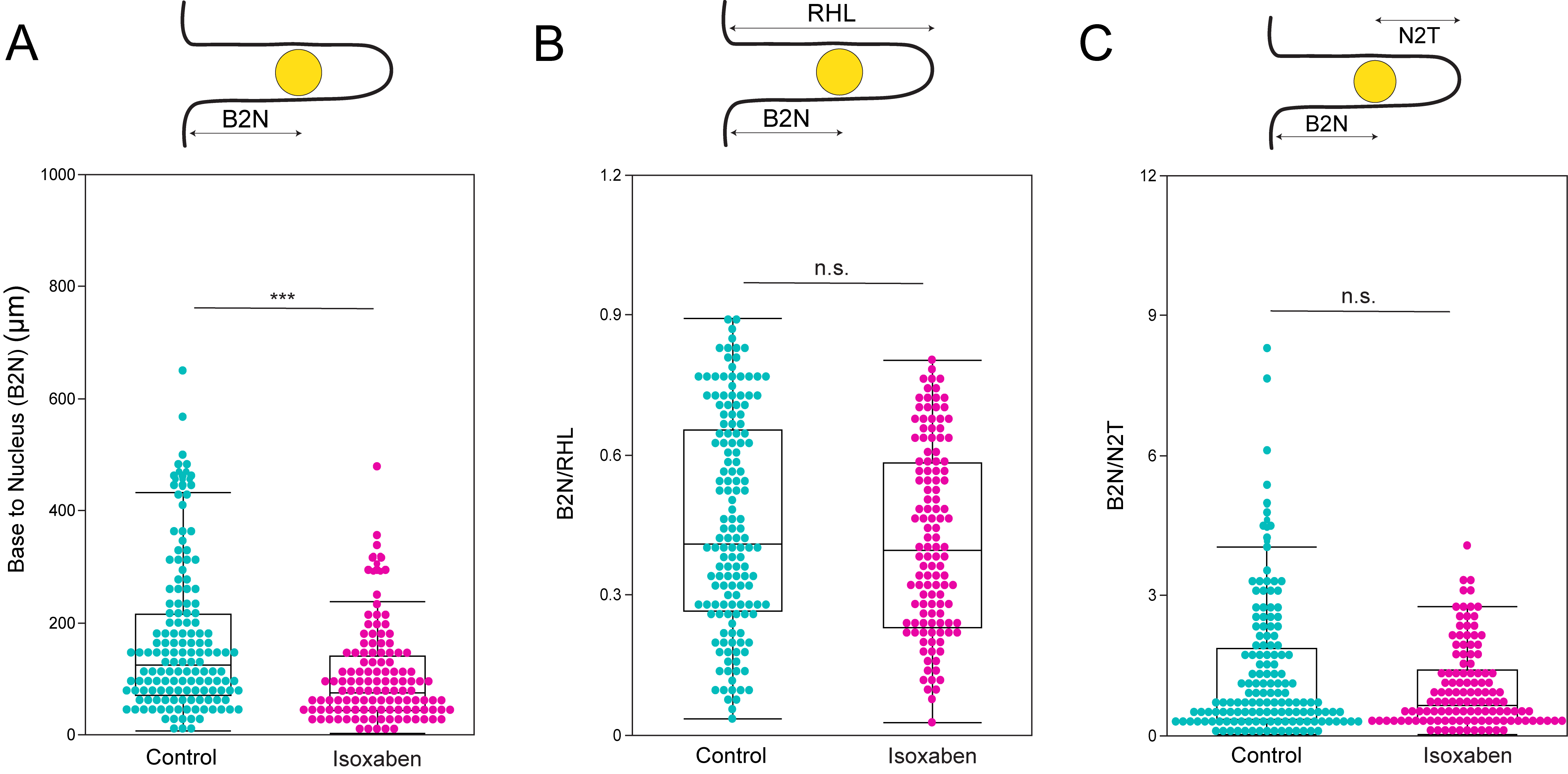

### Supplemental Figure 8.tif

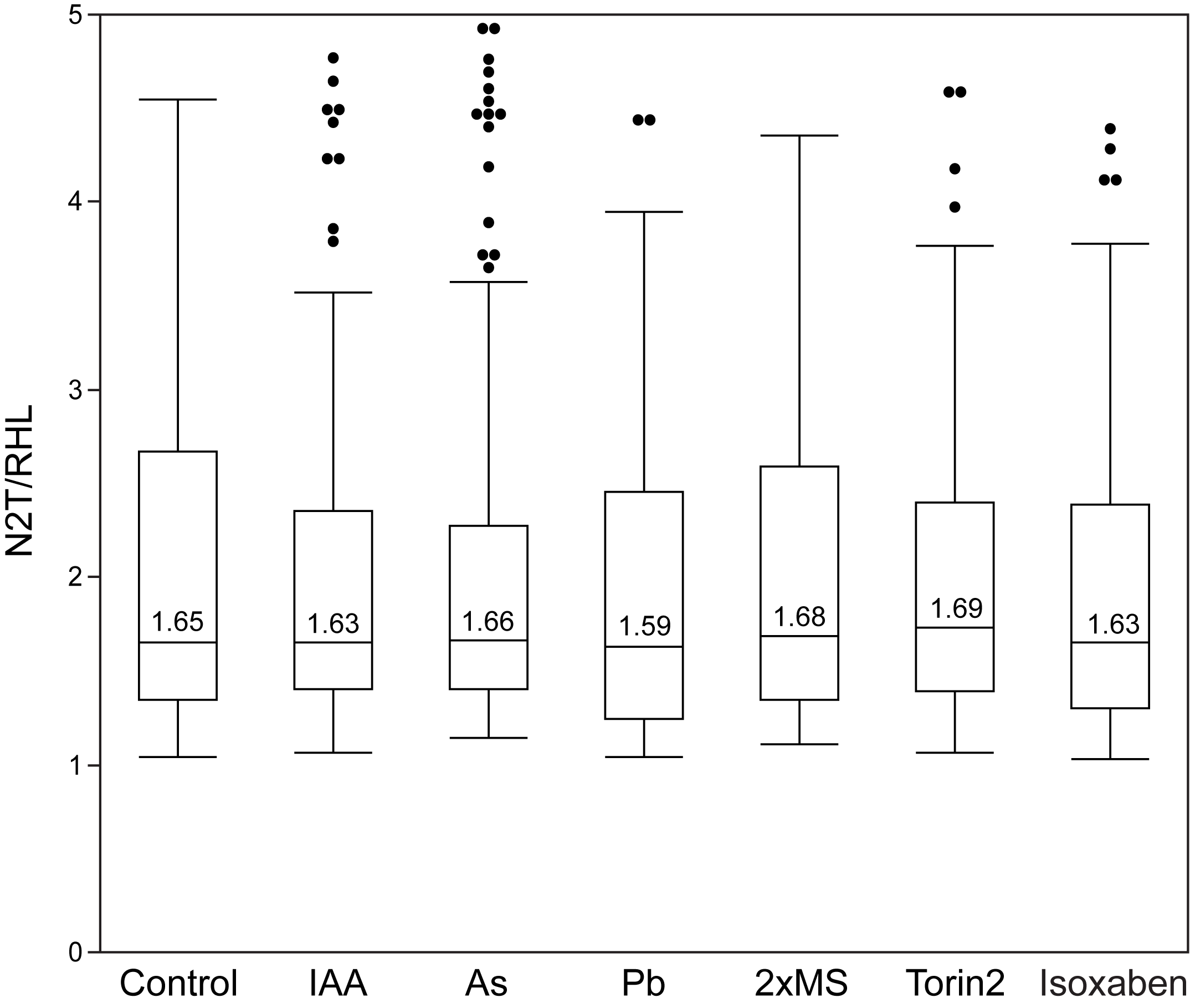
